# A Multi-Objective Genetic Algorithm to Find Active Modules in Multiplex Biological Networks

**DOI:** 10.1101/2020.05.25.114215

**Authors:** Elva-María Novoa-del-Toro, Efrén Mezura-Montes, Matthieu Vignes, Frédérique Magdinier, Laurent Tichit, Anaïs Baudot

## Abstract

The identification of subnetworks of interest - or active modules - by integrating biological networks with molecular profiles is a key resource to inform on the processes perturbed in different cellular conditions. We here propose MOGAMUN, a Multi-Objective Genetic Algorithm to identify active modules in multiplex biological networks. MOGAMUN optimizes both the density of interactions and the scores of the nodes (e.g., their differential expression).

We compare MOGAMUN with state-of-the-art methods, representative of different algorithms dedicated to the identification of active modules in single networks. MOGAMUN identifies dense and high-scoring modules that are also easier to interpret. In addition, to our knowledge, MOGAMUN is the first method able to use multiplex networks. Multiplex networks are composed of different layers of physical and functional relationships between genes and proteins. Each layer is associated to its own meaning, topology, and biases; the multiplex framework allows exploiting this diversity of biological networks.

We applied MOGAMUN to identify cellular processes perturbed in Facio-Scapulo-Humeral muscular Dystrophy, by integrating RNA-seq expression data with a multiplex biological network. We identified different active modules of interest, thereby providing new angles for investigating the pathomechanisms of this disease.

**Availability:** MOGAMUN is available at https://github.com/elvanov/MOGAMUN.

## 1 Introduction

The success of functional genomics is associated with the massive production of quantitative information related to genes, proteins or other macromolecules. These data include, for instance, - omics molecular profiles measuring the expression or activity of thousands of genes/proteins, sensitivity scores resulting from RNA interference or CRISPR screenings, and GWAS scores providing significance of association between genes and phenotypic traits. These scores and measurements, often presented as *p*-values, intend to inform on the cellular responses associated to different cellular contexts. But transforming lists of deregulated genes/proteins and their associated *p*-values to sets of pathways and processes affected in the different cellular conditions remains a major challenge.

A classical approach to identify perturbed cellular processes is the search for over-representation of function or process annotations. Many tools exist that can take as input a list of genes, selected after defining a threshold for significance or ranked according to their *p*-values [1]. Such enrichment approaches will consider only the genes/proteins annotated in databases. Another set of successful approaches try to overcome this limitation by integrating scores or measures with biological networks. Biological networks are composed of nodes representing the biological macromolecules, often genes or proteins, and edges representing physical or functional interactions between those macromolecules. The goal is to identify *active modules*, i.e., subnetworks enriched in interactions and in nodes of interest. These active modules then facilitate the investigation of the perturbed cellular responses, as functional modules are the building blocks of cellular processes and pathways [2].

The identification of active modules from networks is an NP-hard problem [3, 4, 5]. Some active module identification algorithms are based on clustering co-expression networks [6, 7, 8] or memetic algorithms [4]. However, most approaches rely on greedy searches, simulated annealing, and genetic algorithms (see [2] and [9] for general surveys of active module identification methods).

Algorithms based on greedy searches, such as PinnacleZ [10] and MATISSE [11], follow three general steps: i) selection of seed(s), ii) expansion of seed(s), and iii) significance test. In the selection of seed(s), a set of genes of interest (for instance, significantly differentially expressed genes) are picked. Then, the seed(s) are iteratively expanded (adding one node at a time), following a greedy criteria, i.e. choosing the node in the network neighborhood of the seed(s) that maximizes a score, which improves the module fitness. The expansion stops when any of the following three conditions is met: 1) the improvement of the score of the subnetwork is below a minimum threshold, 2) the subnetwork reached a maximum size, or 3) a maximum distance from the seed(s) is reached. As a last step, the subnetworks are tested for significance, by comparing the score of each subnetwork with the score of a random subnetwork. These three steps are common to greedy searches algorithms, but every method has variations. For instance, the seeds selected by PinnacleZ are single nodes, whereas MATISSE selects connected subnetworks. The main drawback of greedy searches is that they can get trapped in local optima because at every step they only look at the local options. In particular, they cannot pick low scoring nodes, even if these can be key for escaping local optima and have access to several high scoring nodes.

Methods based on simulated annealing, such as jActiveModules [3], follow a hill-climbing philosophy, but instead of always picking the best option, i.e., the best neighbor node to be added to the subnetwork, they can also choose unfavorable options (i.e. options decreasing the global score), and thereby escape local optima. Algorithms based on simulated annealing follow two steps: i) initialization of nodes states, and ii) toggling of nodes states. In the initialization of nodes states, each node in the network gets either the active or inactive state, with a given probability. The set of active nodes constitutes the initial subnetwork, and the subnetwork’s score is calculated as the aggregated score of its nodes. Then, in the second step, the nodes states are toggled: in every iteration, the state of a random node is changed from active to inactive, or vice versa. If the toggling improves the score of the subnetwork, it is always accepted; otherwise, it is accepted with a probability calculated based on the temperature parameter, which decreases gradually in every iteration. After toggling states for a given number of iterations, the highest scoring subnetwork found in any iteration is given as result. Some algorithms, such as jActiveModules, have a third step to evaluate the significance of the final subnetwork by comparing its score to scores obtained on randomized expression data. The main drawback of simulated annealing is that the bigger the input network is, the more iterations are needed in order to explore the full search space. Moreover, simulated annealing does not guarantee that the final set of nodes forms a single connected component. However, jActiveModules can filter such set of nodes, in order to keep the top-scoring single connected component(s).

Methods such as COSINE [12], the algorithm proposed by Muraro et al. [13], the one proposed by Ozisik et al. [14] or the one proposed by Chen et al. [5], are all based on genetic algorithms. A key feature of genetic algorithms is that several potential solutions are considered simultaneously. In a genetic algorithm, an initial population of individuals, i.e. subnetworks corresponding to potential solutions, is usually randomly generated. Each individual’s fitness is then evaluated using one (mono-objective optimization) or several (multi-objective optimization) objective functions. The population of individuals then starts the evolution process, where new individuals are generated by crossing existing ones and by modifying them with mutations. The fittest individuals (those with better values for the objective function(s)) have a higher probability to be selected for the generation of offspring. The evolution stops when the algorithm converges, for instance, when there is no improvement in the best value for the objective function(s) for a given number of generations. One of the main advantages of genetic algorithms is that the crossover and mutation operators can help to find a balance between exploring different areas of the whole search space and exploiting the surroundings of promising regions. However, as in simulated annealing, standard crossover and mutation operators cannot guarantee that the final solution will have a set of nodes forming a single connected component. As an option, one can design customized crossover and mutation operators, as in [13, 14]. Importantly, genetic algorithms are capable of optimizing multiple (often conflicting) objectives simultaneously. If the problem is tackled as mono-objective, all the objectives are added into a single objective function by considering weights for each one of them, and the result is usually a single solution. In contrast, if the problem is defined as multi-objective, each objective is associated with an independent objective function, and the result generally leads to several solutions that provide a trade-off for the values of the different objective functions.

By definition, active modules are expected to be enriched in interactions. However, to our knowledge, only few methods, such as SigMod [15], consider the density of interactions. Moreover, existing methods were designed for the analysis of single biological networks, usually a protein-protein interaction network. However, we now have access to several sources of physical and functional interactions between biological molecules. These interactions are represented in a diversity of biological networks, from networks encompassing metabolic and signaling pathways to networks representing correlation of expression. These different interaction networks, each having their own features, topology and biases, are better represented as multiplex networks. Multiplex networks are multilayer networks (i.e., networks composed of different layers, where every layer is an independent network), sharing the same set of nodes, but different types of edges [16]. We and others recently developed different approaches to study and leverage these more complex but richer biological networks [17, 18, 19, 20, 21]. In this work, we present MOGAMUN, a multiobjective genetic algorithm able to explore a multiplex network to identify several active modules.

## 2 Materials and methods

### 2.1 The MOGAMUN algorithm

A multiplex network is defined as a triplet 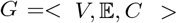, where *V* is the set of nodes, 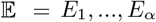 correspond to the *α* different types of edges between the nodes in *V*, one type per layer of the multiplex network, and *C* = {(*v*, *v*, *l*, *k*): *v* ∈ *V*, *l*, *k* ∈ [1,*α*], *l* ≠ *k*} is the set of coupling links that link every node *v* with itself across the *α* layers. For every type of edge in a layer *l*, *E_l_* = {(*v_i_*, *v_j_*): *i* ≠ *j*, *v_i_*, *v_j_* ∈ *V*} [22].

We introduce **MOGAMUN**, a Multi-Objective Genetic Algorithm to identify active modules from MUltiplex Networks. MOGAMUN is a customized version of the Non-dominated Sorting Genetic Algorithm II (NSGA-II) [23], adapted to deal with networks. Our goal is to identify subnetworks that jointly fulfil two objectives: the relevance of the nodes and the density of interactions, inside a given subnetwork. We measure the relevance of the nodes in a subnetwork, using the **first objective function**, the average nodes score, defined in Equation (1).

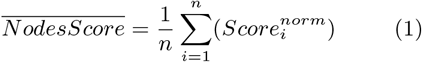

Where *n* is the number of nodes in the subnetwork, and *Score_i_* = Φ^−1^(1 – *p_i_*) is the weight of node *i*. Φ^−1^ is the inverse standard normal cumulative distribution function and *p_i_* is the resulting *p*-value, or FDR-corrected *p*-value, of a statistical test. A node is considered significant if its *p*-value/FDR is lower than a user-defined threshold. In many cases, it corresponds to the result of a differential expression analysis.

The calculus of the inverse normal cumulative distribution (Φ^−1^) leads to values in the range between (–∞, +∞). We use Equation (2) to normalize the nodes scores to be in the [0, 1] range. The average nodes score 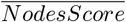 is thus also within this range. Notice that the average nodes score is not an aggregated z-score, as defined in [3], because our 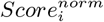 can be computed from either *p*-value or FDRs and is scaled to the range from 0 to 1. It is thereby not necessarily distributed according to a standard normal distribution.

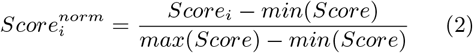

The **second objective function** intends to evaluate the density of interactions in a subnetwork. Here, we compute a normalized density in order to evaluate the density of a subnetwork in a multiplex network. We define the normalized density *D_norm_* in Equation (3).

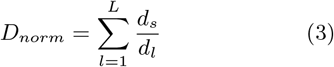

Where *L* is the total number of layers in the multiplex network, *d_l_* is the overall density of layer *l*, and *d_s_* is the density of the subnetwork in layer *l*; the densities *d_s_* and *d_l_* are defined by Equation (4).

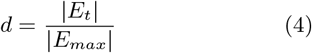

Where *E_t_* is the total number of edges in *d_s_* or *d_l_*, and *E_max_* is the number of edges of the complete graph of the corresponding size.

We present the general flowchart of MOGAMUN in Figure 1. We initialise the algorithm with a random population of individuals (parents). We then mate the initial population to create a new population (children) of same size. Last, we select the best individuals out of the two populations (parents & children) to use them as parents in the next generation. We iteratively repeat the process until convergence. The step-by-step procedure is detailed below, in subsections 2.1.1 to 2.1.10. The algorithm parameters are presented in section 2.2.

**Figure 1:**
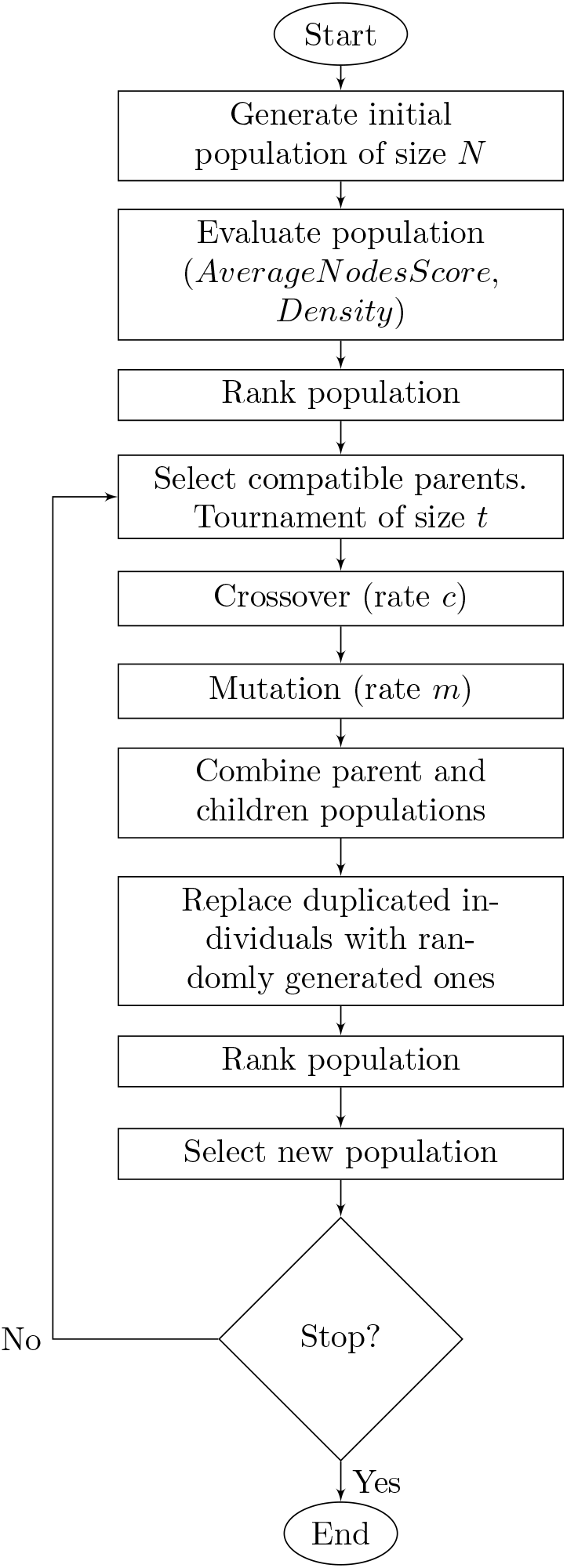
General flowchart of MOGAMUN

We modified NSGA-II [23] to work with networks. To do so, we defined a coding scheme for the individuals with a variable length, where each feature corresponds to the identifier of a node. We also customized the original steps involving either the creation or the modification of individuals (generation of the initial population, crossover and mutation). In addition, we added a step to replace duplicated individuals with randomly generated ones, in order to ensure the diversity of the population and allow exploring the search space further. Importantly, we request all the individuals (i.e., the subnetworks of the multiplex network) to be single connected components.

#### 2.1.1 Generating the initial population

We first defined a multiplex-network version of the Depth First Search (multiplex-DFS, see Algorithm 1), which allows generating individuals that are single connected components. In every iteration of the multiplex-DFS, a uniformly random layer of the multiplex network is visited (see Algorithm 1, line 9). We use the multiplex-DFS to generate an initial population of *N* individuals. Each individual is a connected subnetwork with a random size between *MinS* and *MaxS*. The seed, i.e., the initial node in the network, is randomly chosen from the pool of significant nodes, in order to focus around interesting areas of the multiplex network.

**Algorithm 1.**
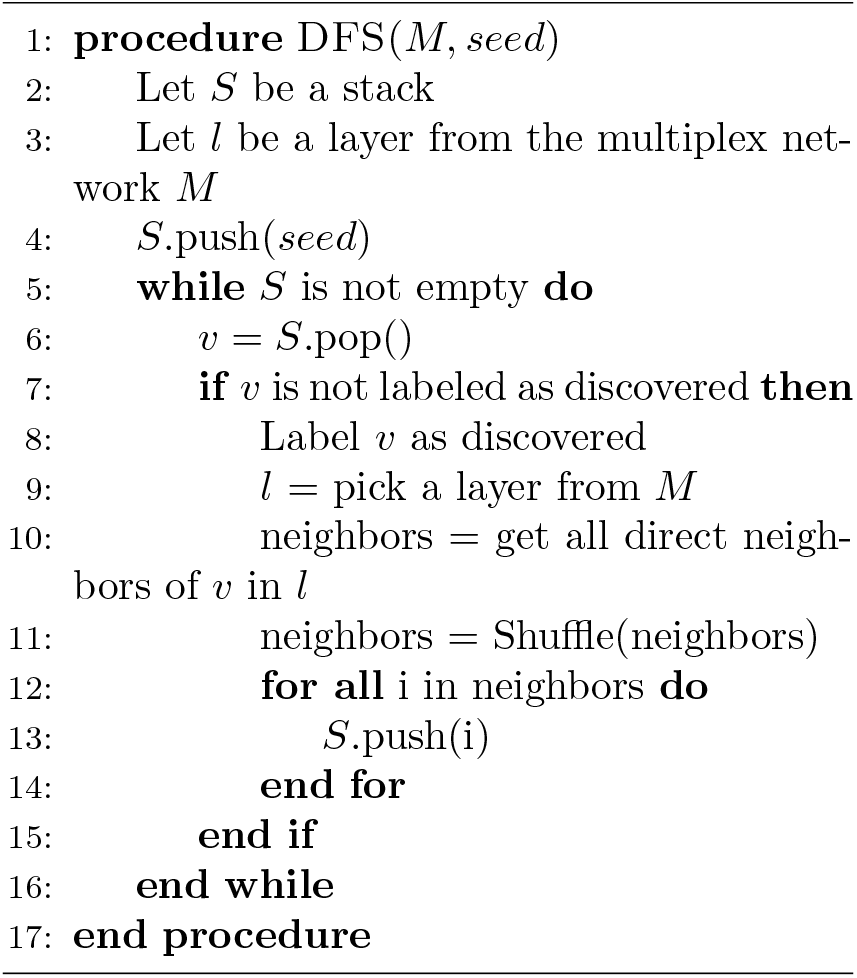
Multiplex Depth First Search

#### 2.1.2 Evaluating the initial population

We now evaluate all the individuals of the population, i.e., the set of potential subnetwork solutions, with the two objective functions described in Equations (1) and (3). A high average nodes score implies that the individual contains high-scoring nodes. Similarly, a high normalized density implies that the individual is densely connected in the multiplex network.

#### 2.1.3 Ranking individuals in the population

We use the Pareto dominance, a classical criterion in evolutionary multi-objective optimization, to rank the individuals [24]. In a maximization problem, an individual *S*1 dominates *S*2 (*S*1 ≻ *S*2), if 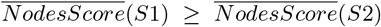 and *D_norm_*(*S*1) ≥ *D_norm_*(*S*2), and at least one of the two inequalities is strict. The ranking process is carried out like in the original NSGA-II algorithm, as follows: initially, all non-dominated individuals (i.e., those individuals that are not dominated by any other individual in the current population) are assigned rank 1 and separated from the population. After that, from the remaining individuals in the population, those non-dominated are assigned ranking 2 and separated from the population as well. Such process continues until there are no remaining individuals in the current population. At the end, all individuals in the population have a ranking value. The best individuals have rank 1.

Apart from assigning a rank to every individual, we also calculate their crowding distance, which is a measure that determines the proximity of the individuals in the objective space. The crowding distance of an individual is equivalent to the perimeter of the cuboid formed by its surrounding nearest pair of individuals in the same Pareto front, one at each side. The only exception is for those individuals that maximize one of the two objectives in each rank, which are directly assigned an infinite crowding distance value [24].

#### 2.1.4 Selecting compatible parents by tournament

The parents are selected by tournament [25]. The selection of a pair of parents restricts the crossover to individuals that are compatible. This ensures that the children are also single connected components. Two individuals *S*1 and *S*2 are compatible if:

- 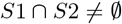, or
- 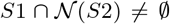, where 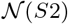 is the set of neighbors of the nodes of *S*2.

The first parent is chosen via tournament (considering the rank of the individuals, and the crowding distance if they have the same rank). Depending on the number of compatible individuals, the second parent can be either selected also by tournament or directly assigned, if there is only one compatible individual. The procedure is described in Algorithm 2. If no individual is compatible with the first parent, we restart the process with a different individual as *Parent*1 (line 10 of Algorithm 2). If after a pre-specified number of attempts, the search of compatible parents is unsuccessful, we add a copy of two individuals (randomly chosen) to the population of children and we skip crossover and mutation.

**Algorithm 2.**
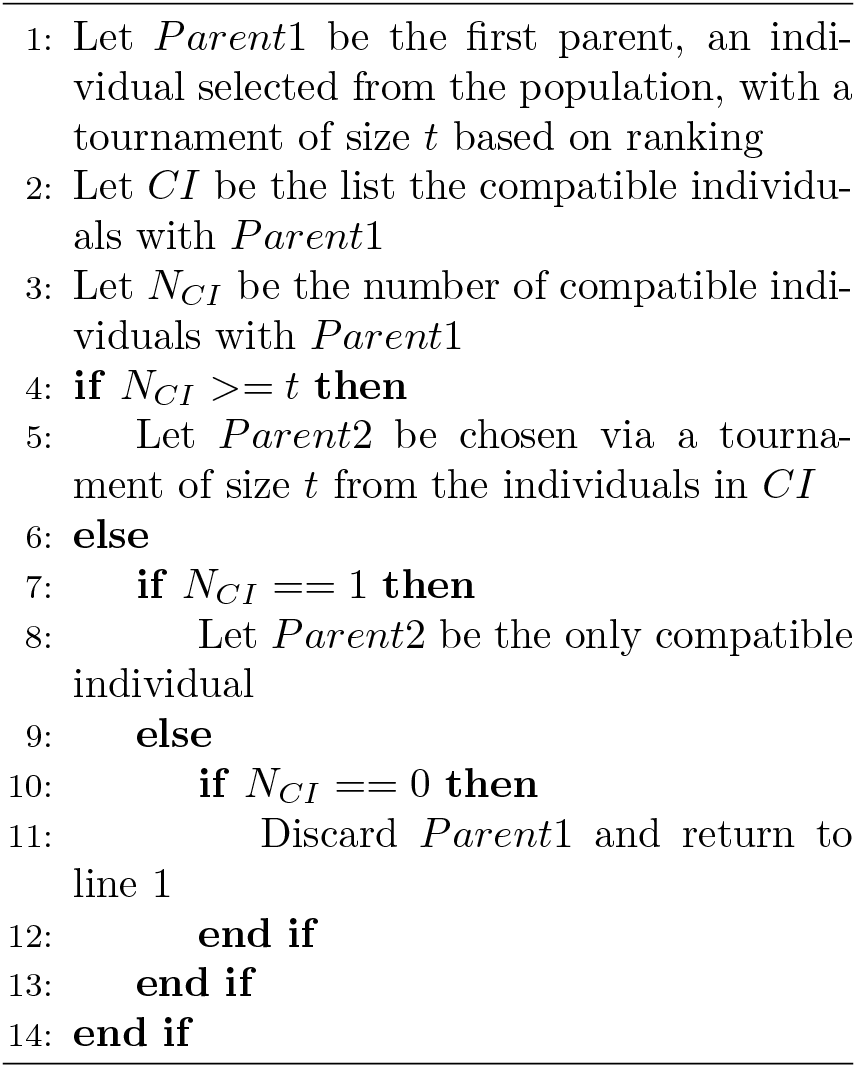
Selection of compatible parents

#### 2.1.5 Crossover

The goal of the crossover operator is to combine the nodes of two parent individuals, in an attempt to improve the values of any of the objective functions (average nodes score or normalized density). We mate the parents with crossover rate *c*. In order to guarantee that each child will be a single connected component, we use a crossover method inspired from the one proposed in Muraro et al., where the subnetworks corresponding to the parents are merged to have a single connected component [13]. In such a way, two nodes are randomly chosen, and two new children are generated with a Depth First Search, having as seed each selected node, respectively. However, our crossover varies according to two main aspects: 1) each seed for the children must correspond to significant nodes, and 2) the children can be generated either with Depth First Search or Breadth First Search. All children respect the subnetwork size’s range.

#### 2.1.6 Mutation

The goal of the mutation operator is to exploit the neighborhood of the children, adding/removing nodes, here also in an attempt to improve the value of any of the two objective functions. Notice that a node that is in the neighborhood of a child, i.e., directly connected to it, and that has a high node score, would allow increasing the average nodes score. In the same way, a neighbor node that is highly connected with the nodes of the child could improve the normalized density. We mutate each child independently with rate *m*. We first choose the list of potential nodes *v_p_* to be removed. We restrict this list to those nodes that can be removed without disconnecting the child subnetwork and that are not significant. We finalize the mutation process by adding |*v_p_*| new nodes to the subnetwork if |*v_p_*| > 0, or a single node if |*v_p_*| = 0. The new nodes are chosen randomly, from the neighborhood of the corresponding child, considering all the layers of the multiplex network, and preferring significant nodes, if existing.

#### 2.1.7 Combining parent and children populations

We join the parent and children populations, giving as result a population of size 2*N*.

#### 2.1.8 Replacing duplicated individuals with randomly generated ones

Duplicates of individuals appear in the population when no compatible parents are found or when no crossover nor mutation are applied. To preserve diversity in the population, promote the exploration of the search space and avoid premature convergence, we introduce the replacement of duplicated individuals. To determine if an individual is duplicated, we check if it has more nodes in common with another individual than a given threshold *J_t_* of the Jaccard similarity coefficient. The Jaccard similarity coefficient *J* between two subnetworks (individuals) *A* and *B*, is calculated as follows:

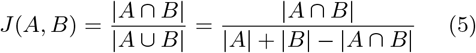

The dominated individual is labeled as duplicated. If no domination exists, one of the individuals is randomly selected.

#### 2.1.9 Selecting the new population

After ranking the full population of size 2*N*, the new population of size *N* is selected with elitism. The top *N* ranked individuals will form the new population, whereas the other *N* are discarded. It is to note that the best individuals (among the parents and children) will thereby always remain in the population.

#### 2.1.10 Stopping criteria

At this point, we have completed one generation. We iterate until reaching the stopping criteria, given by the number of generations *gen*. The result is the set of individuals in the **first Pareto front** (rank = 1). Evolutionary algorithms are stochastic search approaches, they must hence be run several times. As a result, we will have several first Pareto fronts. In order to select the final set of individuals, we calculate the *accumulated Pareto front*. To this goal, we take the results of all runs, re-rank them, and keep only those individuals in the new first Pareto front.

### 2.2 Parameter values

In the study presented here, we used the parameter values listed in Table 1. We generated subnetworks in size range of [15–50], which corresponds to the size of communities identified by four over five top-performing algorithms in a community identification challenge [26]. This size range combined with a population of size 100 individuals, allows covering about a quarter of the multiplex network, around the most interesting areas. Tournament size and crossover rate are classical values in genetic algorithms. Mutation rate of 10% is higher than in most approaches, to promote the exploitation of the search space near good solutions. We elected the total number of generations empirically, after running the algorithm several times in different contexts and monitoring its convergence. Similarly, the threshold of the Jaccard similarity coefficient was also obtained empirically; we found that it allows to keep a high diversity rate, while preventing pre-mature convergence.

**Table 1:** Parameters of MOGAMUN and values used in this study.

| Feature | Description | Value |
| --- | --- | --- |
| $N$ | Population size | 100 |
| $MinS$ | Minimum size of the individuals | 15 |
| $MaxS$ | Maximum size of the individuals | 50 |
| $t$ | Tournament size | 2 |
| $c$ | Crossover rate | 80% |
| $m$ | Mutation rate | 10% |
| $gen$ | Total number of generations | 500 |
| $J_t$ | Jaccard similarity coefficient threshold | 30% |

### 2.3 Benchmark to compare the performance of MOGAMUN with existing methods

In order to compare the performance of MOGAMUN with state-of-the-art approaches, we used and extended the benchmark initially proposed by Batra et al. [27]. It is worth noticing that this benchmark works with a single interaction network as, to our knowledge, no active module identification method can consider multiplex networks. It artificially generates expression data to simulate a differentially expressed subnetwork. To this goal, the nodes of the network (i.e., the genes) are separated into two groups: Foreground Genes (FG) and Background Genes (BG). A *seed-and-select* algorithm is defined to randomly select the FG as a connected subnetwork. Artificial expression data is then generated so that the FG contrasts with the BG, which means that the FG genes are artificially differentially expressed.

We computed the *F*_1_ score (also known as *F* score or *F* measure) to evaluate the quality of the active modules identified by the different methods. The *F*_1_ score is calculated on the union of all the active modules retrieved by each method over the 30 runs, using the equation 6.

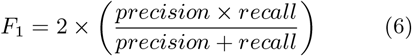

Where 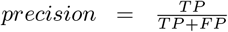 and 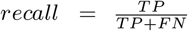. The *TP* and the *FP* are the number of FG and BG present in the active modules, respectively, and the *FN* are the number of missing FG (i.e. the FG that were not retrieved).

#### 2.3.1 Benchmark Networks

We used two independent interaction networks in the benchmark (Table 2). *PPI*_1 is the human protein reference database [28], taken from [27]. *PPI*_2 was generated by merging interactions identified from several databases through the PSICQUIC portal [29] and the CCSB Interactome database [30], taken from [17].

**Table 2:** Interaction networks used in the benchmark.

| Name | Number of nodes $\vert V\vert $ | Number of edges $\vert E\vert $ | Density $d$ |
| --- | --- | --- | --- |
| *PPI_1* | 9 425 | 36 811 | $8.28 \times 10^{-4}$ |
| *PPI_2* | 12 621 | 66 971 | $8.41 \times 10^{-4}$ |

#### 2.3.2 Benchmark artificial expression data

We simulated 2 different artificial expression datasets, one following a normal distribution and another one sampled from real RNA-seq data, as follows:

1. *Simjnormal*. We simulated data following a normal distribution. The mean values *μ* of the FG and BG groups of genes are *μ*(*FG*) = 5, and *μ*(*BG*) = 2, respectively, and a standard deviation SD = 1 for both groups of genes. This situation corresponds to a high signal strength, as in [27]. To test for differential expression, we performed a series of *t*-tests, and considered a gene as significantly differentially expressed if the *p*-value ≤ 0.05. We used a set of 20 FG.
2. *Samp_TCGA*. We sampled data from real expression data, in order to have an RNA-seq distribution-like. We downloaded breast cancer RNA-seq expression dataset from the TCGA breast cancer project (TCGA-BRCA) from the US National Cancer GDC portal (https://portal.gdc.cancer.gov/), as of May, 2019. This dataset is composed of 1 102 cases and 112 controls (we removed the outlier control sample “d5f0ea64.6660.49ac.a37e.3cd747045147”). To test for differential expression, we used the R package *edgeR* version 3.26.8 [31]. We consider a gene to be significantly differentially expressed if its False Discovery Rate *FDR* ≤ 0.05 and |*log*_2_(*FC*)| > 1, where the FC is the ratio of the difference in expression between cases/patients and controls. The expression data for the FG and BG were randomly sampled from the set of significantly differentially expressed and non-differentially expressed genes, respectively. We used a set of 20 FG.

In Table 3 we describe our two datasets. Columns “Cases” and “Controls” show the number of patients/cases and controls, respectively, “Genes” shows the total number of genes in the simulated dataset, corresponding to the total number of nodes in the networks, and “Significant DE” is the number of significantly differentially expressed genes.

**Table 3:** Artificial expression datasets.

| Name | Cases | Controls | Genes | Significant DE |
| --- | --- | --- | --- | --- |
| *Sim_normal* | 100 | 10 | 9 425 | 483 |
| *Samp_TCGA* | 1 102 | 112 | 12 621 | 20 |

#### 2.3.3 State-of-the-art algorithms selected for comparison

We compared MOGAMUN with three selected methods, representative of the main approaches seeking for active modules: jActiveModules [3], PinnacleZ [10], and COSINE [12].

##### 2.3.3.1 jActiveModules

Ideker et al. [3] proposed jActiveModules. They use a simulated annealing algorithm to find subnetworks with the highest scores, calculated from the differential expression of the subnetwork nodes. The search starts by selecting a subnetwork containing approximately half of the nodes of the full network. After that, they iteratively add or remove one node at a time from the selected subnetwork (the number of iterations is defined *a priori*). Whenever the addition or removal of a node increases the score of the subnetwork, the modification is accepted. Otherwise, it is accepted with a probability that decreases along the iterations, according to the temperature value. After finishing adding/removing nodes, the highest scoring subnetwork (found in any iteration) is selected as result. Finally, the significance of the selected subnetwork is evaluated. Several parameters can be tuned, but for the tests performed here, we used the default values recommended by the authors [3]. The only exception is that we set to 1 the number of modules to be retrieved, as this corresponds to the benchmark settings. jActiveModules is available as a Cytoscape plugin.

##### 2.3.3.2 COSINE

Ma et al. [12] proposed COSINE, a method based on a standard mono-objective genetic algorithm. The goal of COSINE is to find the subnetwork with the highest change in expression among conditions, represented as node weights. COSINE further allows considering the level of co-expression between pairs of genes, represented as edge weights. To compute the edge weights, we calculated the coexpression of every pair of nodes connected in the benchmark networks. COSINE further allows giving more importance to either the weights of the nodes or the edges, with a parameter lambda. For the tests performed here, we used the same parameters as reported in [12], where COSINE is compared with other methods (number of iterations = 5,000; zero to one ratio = 30), and we set lambda to 0.5, in order to give the same importance to changes in expression (i.e. node weights) and co-expressions (i.e. edge weights). COSINE is available as an R-package.

##### 2.3.3.3 PinnacleZ

Chuang et al. [10] designed PinnacleZ, a greedy algorithm to identify active subnetworks that maximize the mutual information. The mutual information measures the differences in the distribution of the expression values of a given set of genes between two conditions. PinnacleZ starts the search by selecting an initial set of seeds, and for each of these seeds, it iteratively adds the neighbor node that maximizes the mutual information of the subnetwork. The search stops when a maximal distance from the seed is reached or when the improvement of the mutual information score is not considered significant, given a threshold. PinnacleZ then performs three tests of significance on each of the identified active subnetworks, in order to guarantee that their individual mutual information is higher than the mutual information of a random subnetwork. We used the same parameters reported in [10] (distance from the seed = 2 nodes, minimal mutual information score improvement threshold = 0.05), and we set the maximum size per subnetwork = 50 (the same size that we allowed for MOGAMUN). PinnacleZ was originally available as a Java program and a Cytoscape plugin, but this latter one is no longer supported.

### 2.4 Application to Facio-Scapulo-Humeral muscular Dystrophy type 1 (FSHD1)

#### 2.4.1 RNA-seq expression data

We used five Facio-Scapulo-Humeral muscular Dystrophy type 1 (FSHD1) RNA-sequencing expression datasets publicly available [32, 33, 34], extracted from the Gene Expression Omnibus [35]. We performed the differential expression analyses using the R package *edgeR* version 3.26.8 [31]. As recommended in the user guide of *edgeR*, we performed glmQLF tests for the two datasets with samples from different batches [33, 34], and Fisher Exact tests for the three datasets with samples from a single batch [32]. We considered a gene as significantly Differentially Expressed (significantly DEG) if the False Discovery Rate *FDR* ≤ 0.05 and the |*log*_2_(*FC*)| > 1, where the FC (Fold-Change) is the ratio of the difference in expression between cases and controls.

We selected from Yao et al. [32] RNA-seq data from muscle biopsies of 9 FSHD1 patients (quadriceps, 4 males and 5 females) and 9 controls (8 quadriceps, 1 tibialis anterior, 5 males and 4 females). We also selected data from two myoblasts derived from patients and the two corresponding myotubes, as well as two myoblasts from controls and three control myotubes (Supplementary Table S1). The cells were obtained from the University of Rochester repository and are described in Young et al. [36]. Our differential expression analyses revealed 6, 7 and 343 significantly DEGs, for biopsies, myoblasts and myotubes, respectively.

In Banerji et al. 2017 [33] RNA-sequencing was performed in triplicate on confluent immortalized myoblasts, for three FSHD1 patients (corresponding to 5 cell lines, for a total of 15 samples) and three healthy individuals (corresponding to 4 cell lines, for a total of 12 samples) (Supplementary Table S2). These cells were on one hand derived from a mosaic patient and described in Krom et al. [37] (54-12; 54-45; 54-2 for FSHD1 cells with 3 D4Z4 units and 54-6; 54-A10 as controls with 13 D4Z4 units). On the other hand, the 12Ubic and 16Ubic cells obtained from two FSHD1 patients and the 12Abic and 16ABic cells from matching controls are described in Homma et al. [38]. We identified 192 significantly DEGs comparing all the FSHD1 to control samples.

The last dataset was obtained from Banerji et al. 2019 [34] and corresponds to myotubes collected at the end of myoblasts to myotubes differentiation. These myotubes are derived from the myoblasts described in Banerji et al. 2017 [33]. In Banerji et al. 2019, a time course expression during differentiation was analyzed. We considered here only the last time point (T8) and selected triplicated samples for 5 FHSD1 patients and 4 controls (Supplementary Table S3). We identified 261 significantly DEGs.

#### 2.4.2 Biological interaction networks

We built a multiplex network composed of three layers of physical and functional interactions (see Table 4). The nodes are either genes or proteins, considered here equally. The edges are undirected, and we removed loops (i.e., self-interactions). The three networks were taken from [17]. The first network (*PPI_2*), is a protein-protein interaction network, and is also the one used for the benchmark (Section 2.3). In the second network (*Pathways*), the links correspond to pathway interaction data, obtained with the R package *graphite* [39]. The last network (*Co-expression*), contains edges corresponding to correlations of expression. Spearman correlations were calculated from RNA-seq data of 32 tissues and 45 cell lines, and absolute correlations of at least 0.7 were selected to build the network [40].

**Table 4:**
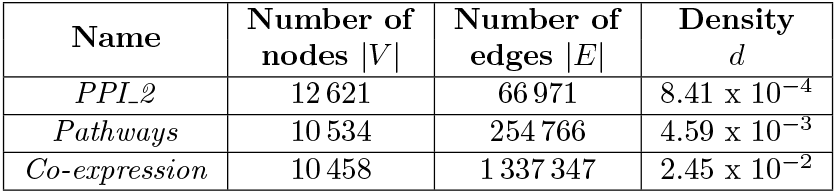
Multiplex biological network.

## 3 Results

To the best of our knowledge, MOGAMUN is the first algorithm that detects active modules from multiplex networks. However, several methods exist to detect one [4, 11, 12], or several [3, 5, 6, 7, 10, 13] active modules in monoplex networks -aka single networks. We here compare MOGAMUN with three state-of-the-art approaches to detect active modules in monoplex networks (section 3.1). We then applied our algorithm to study Facio-Scapulo-Humeral muscular Dystrophy type 1 (FSHD1), using a multiplex network (section 3.2).

### 3.1 MOGAMUN against state-of-the-art active module identification methods

We ran jActiveModules, COSINE, PinnacleZ and MOGAMUN 30 times (see Materials and Methods). The execution times per run of each algorithm, in a desk computer with Intel processor i7 at 3.60GHz and 32GB of RAM, were approximately 30 min, 8 hours, 30 min, and 12 hours for jActiveModules, COSINE, PinnacleZ and MOGAMUN, respectively.

As a first test, we used the *PPI_1* network (Table 2) and the *Sim_normal* dataset (Table 3) (see Materials and Methods). The goal is to retrieve the active module, which is a single subnetwork composed of 20 nodes (i.e., the foreground genes (FG)). The four methods retrieved the 20 nodes of the FG (Figure 2a). PinnacleZ retrieved 13 231 subnetworks in total, corresponding to 494 subnetworks with at least one different node. These 494 subnetworks have an average size of 6 nodes and 3% average Jaccard similarity between them. COSINE and jActiveModules retrieved 30 subnetworks each, one per run. The 30 subnetworks found by COSINE all have at least one different node, an average size of 640 nodes and 5% average Jaccard similarity. Finally, 29 out of the 30 subnetworks retrieved by jActiveModules have at least one different node, an average size of 6 952 nodes and 76% average Jaccard similarity. MOGAMUN retrieved 6 modules with at least one different node, an average size of 17 nodes and 13% average Jaccard similarity. We calculated the *F*_1_ score (Material and Methods) of the union of all the active modules retrieved by each method on the 30 runs. the *F*_1_ score determines how good the methods are to retrieve the foreground genes (FG) while avoiding picking background genes (BG). The *F*_1_ score for jActiveModules, COSINE and PinnacleZ is near to zero (< 1), and > 4 for MOGAMUN (Supplementary Figure S9).

**Figure 2:**
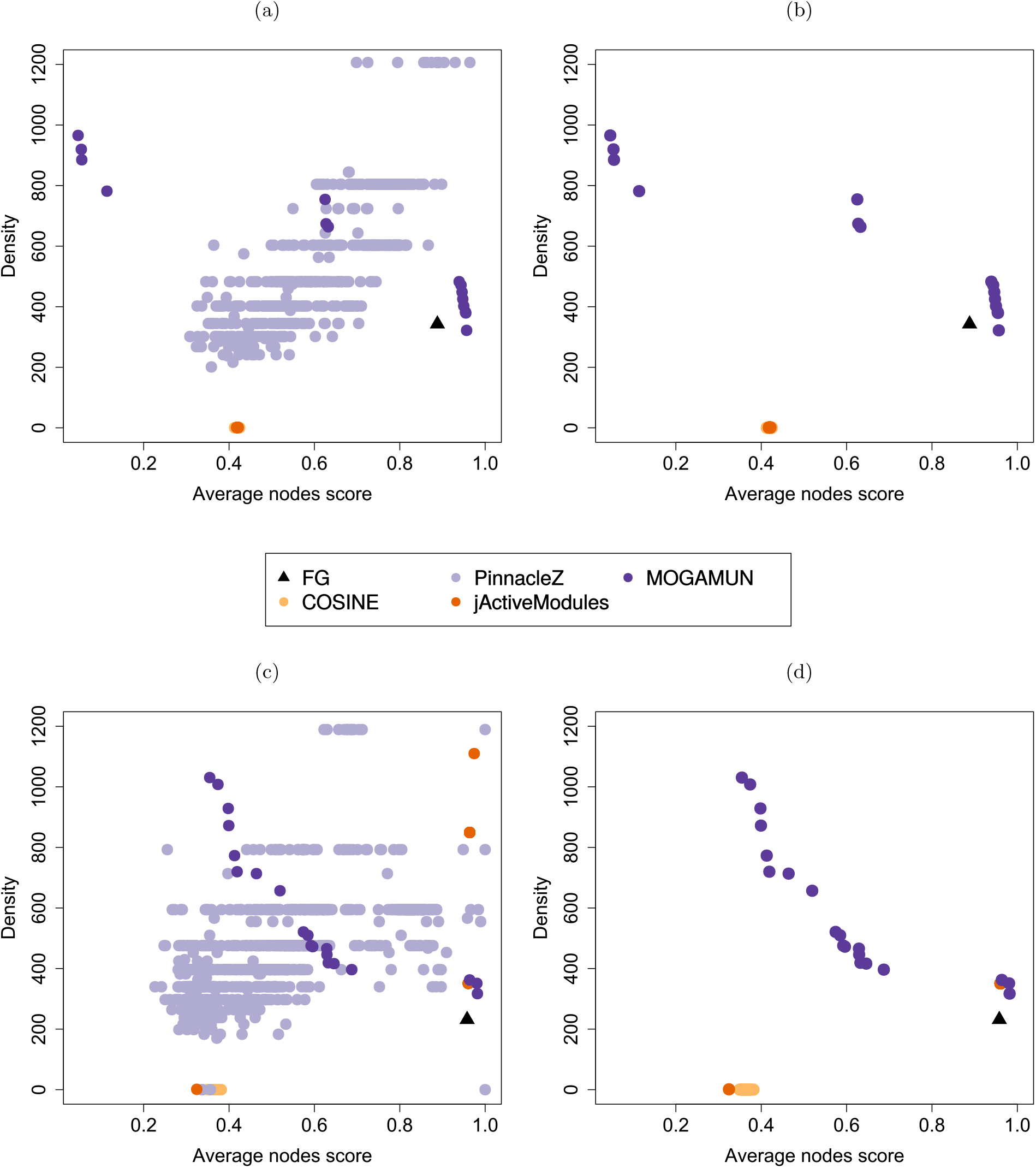
Comparing the performance of MOGAMUN, COSINE, PinnacleZ and jActiveModules. (a) Results of 30 runs using the *PPI_1* network and the *Sim_normal* dataset. (b) Filtered results from (*a*), keeping only the subnetworks with at least 15 nodes. (c) Results of 30 runs using the *PPIJ2* network and the *Samp_TCGA* dataset. (d) Filtered results from (c), keeping only the subnetworks with at least 15 nodes. The size distributions of all the modules can be retrieved in Supplementary Figures S1-8.

Overall, PinnacleZ identified the best modules in terms of both average nodes score and density (figure 2a). However, the identified modules are small (average size of 6 nodes). PinnacleZ indeed restricts the search locally around the seed neighborhood by considering only the subset of nodes that are (at most) two steps away from the seed. In such small modules, the two objectives are expected to reach their maximum values. For example, a subnetwork composed only of two significant nodes with score = 1 linked by an interaction will have a maximal average nodes score, as well as a maximal subnetwork density because it is a complete graph. In order to consider this, we filtered the all the modules obtained by the four methods to keep only the subnetworks with at least 15 nodes (Figure 2b). This removed all the active modules obtained by PinnacleZ, and revealed that MOGAMUN succeeded to find the best results, in terms of the two objectives. MOGAMUN is the only approach allowing setting a minimum allowed size from the four methods implemented here. It is to note that, if we also remove the subnetworks with more than 50 nodes (the maximum size we set in MOGAMUN and PinnacleZ, the two methods allowing this setting), we would also discard all the results from COSINE and jActiveModules, which obtain subnetworks with hundreds or even thousands of nodes.

In a second test, we used the *PPI_2* network (Table 2) and the *Samp_TCGA* dataset (Table 3). The goal is also to retrieve a single active module, which is a subnetwork composed of 20 nodes (i.e., the foreground genes (FG), see Materials and Methods). jActiveModules retrieved the highest number of FG genes (19/20), whereas PinnacleZ and MOGAMUN found 18/20 each, and COSINE, 12/20 (Figure 2c). The results are similar to the ones obtained in the first test. PinnacleZ found 25 313 modules, out of which 1 055 have at least one different node. These 1 055 subnetworks have an average size of 6 nodes and 5% average Jaccard similarity between them. COSINE and jActiveModules retrieved 30 modules each, one per run. The 30 subnetworks found by COSINE all have at least one different node, with an average size of 205 nodes and 5% average Jaccard similarity. Four out of the 30 subnetworks retrieved by jActiveModules have at least one different node, with an average size of 1 033 nodes and 28% average Jaccard similarity between these 4 subnetworks. MOGAMUN retrieved 18 modules with at least one different node, an average size of 16 nodes and 18% average Jaccard similarity. The *F*_1_ score of the union of all the active modules retrieved by each method on the 30 runs is < 1 for jActiveModules, COSINE and PinnacleZ, and > 3 for MOGAMUN (Supplementary Figure S9).

The filtering of the modules having more than 15 nodes in this comparison also removed all the results obtained by PinnacleZ, as well as two high-scoring subnetworks from jActiveModules (figure 2d).

After filtering on module size, MOGAMUN led to the best results, although jActiveModules succeeded to find a module with an average nodes score similar to one of ours. However, MOGAMUN found overall denser subnetworks. If we also remove the subnetworks with more than 50 nodes, we would again discard all the results from COSINE and jActiveModules, with the exception of a single subnetwork, found by jActiveModules. In summary, MOGAMUN clearly identifies the best modules in terms of the multiple objective setting. Moreover, the retrieved modules have reasonable and tunable sizes.

### 3.2 Application to FSHD1

Facio-Scapulo-Humeral muscular Dystrophy type 1 (FSHD1) is a rare autosomal dominant genetic disease characterized by a progressive and asymmetric weakening of specific groups of muscles, with progression from the face to the lower limbs. The particularity of this disease resides in the absence of mutation in a gene encoding a muscle-specific factor. FSHD1 is however associated to a variable number of tandem repeats in the disease locus at the subtelomeric 4q35, more specifically to an array of 3.3 kb macrosatellite elements (D4Z4). In unaffected individuals, this array comprises between 11 and up to an average of 75 units [41]. In patients, this array is shortened with a threshold limit of less than 10 units. D4Z4 encodes the DUX4 transcription factor. The current pathological model associates D4Z4 array shortening with chromatin relaxation, expression of the DUX4 transcription factor and subsequent activation of a number of target genes of poorly known function in muscle physiology [42]. Overall, the biological processes leading to the muscle defects remain currently unclear.

We aim here to apply MOGAMUN in order to reveal biological processes that would not have been exposed by previous analyses, and further define biomarkers associated with the muscle phenotype of patients. We applied MOGAMUN using a multiplex networks composed of three layers of biological interactions and FSHD1 RNA-sequencing expression datasets obtained from different types of cells [32, 33, 34] (Materials and Methods). More precisely, the first FSHD1 RNA-seq datasets were obtained from biopsies, myoblasts and myotubes differentiated from those myoblasts [32]. The two other datasets were obtained from immortalized myoblasts [33] and myotubes differentiated from those myoblasts [34] (Material and Methods). We independently ran MOGAMUN 30 times.

We first considered the results obtained from [32] dataset, and analyzed the active modules identified in biopsies (18 active modules, Supplementary Figures S10), myoblasts (10 active modules, Supplementary Figures S11) and corresponding myotubes (23 active modules, Supplementary Figures S12).

In myoblasts, all the 10 active modules contain at least one of the two significantly down-regulated genes LRRTM4 and GFRA1 (Supplementary Figures S11). LRRTM4 is required for presynaptic differentiation and GFRA1 belongs to the GDNF family receptor, also involved in the control of neuron survival and differentiation. The function of these two factors in muscle cells is, to our knowledge, not described. However, it is interesting to note that they belong to active modules containing proteins implicated in ubiquitination, intracellular signaling and DNA replication machinery, as FSHD1 cells display increased apoptosis and reduced proliferation.

Using expression data obtained from myotubes differentiated from these myoblasts, MOGAMUN identified 23 active modules, among which many contain a subset of highly over-expressed nodes (Supplementary Figure S12). These nodes associated to a high fold-change are DUX4 target genes, as defined in Geng et al. [42] and in Yao et al. [32] from DUX4-transduced over-expression experiments. In the active modules, the DUX4 target genes are however only linked together by interactions inferred from correlation of expression, and do not share pathway nor physical interactions. The over-expression of DUX4 target genes in differentiated cells is consistent with previous observations showing increased expression of this gene and its target genes upon differentiation. In some active modules, the DUX4 target genes are connected to cyclins through the Ubiquitin conjugating enzymes E2 D1 (Figure 3a). They are connected in particular to CCNA1, involved in cell cycle regulation at the G1/S and G2/M, and also reported as a DUX4 target gene in DUX4-transduced [42, 32] and immortalized [37, 43] myoblasts.

**Figure 3:**
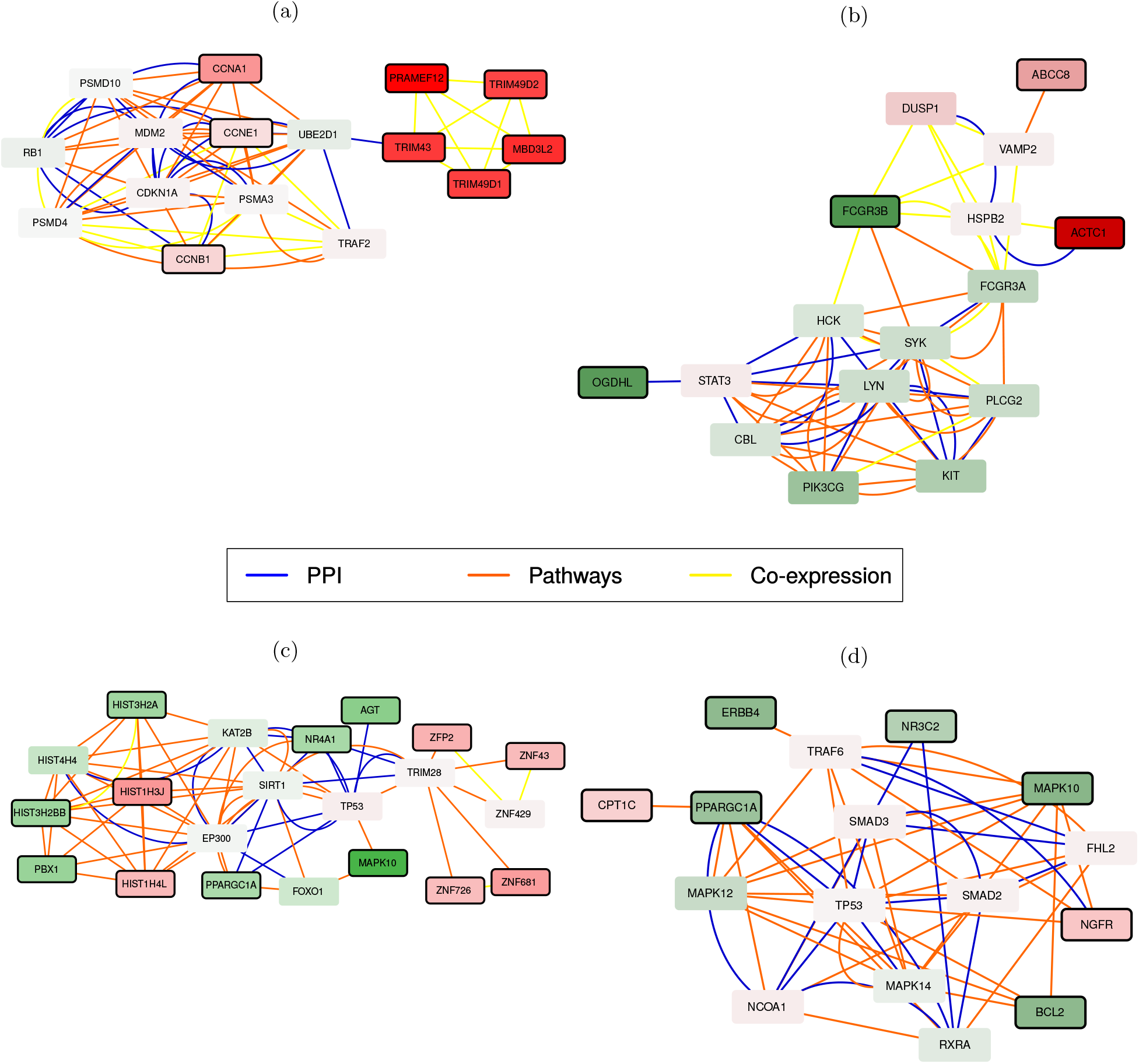
Four active modules obtained by applying MOGAMUN on different FSHD1 expression datasets. The color of the nodes represents the fold-change, where green and red nodes correspond to under- and over-expressed genes, respectively. Nodes with bold black border correspond to genes significantly differentially expressed (*FDR* < 0.05 and absolute *log*_2_ fold-change > 1). Blue nodes correspond to genes with no associated transcriptomics data. The active modules are extracted from the sets of active modules obtained from a) myotubes from Yao et al. 2014 [32], b) biopsies from Yao et al. 2014 [32], c) myoblasts from Banerji et al. 2017 [33] and d) corresponding myotubes from Banerji et al. 2019 [34].

In biopsies, Yao et al. also detected some DUX4 targets genes at a low level [32]. However, we do not identify DUX4 target genes in active modules obtained by MOGAMUN from biopsy expression data. Notably, in biopsies, we identified an interesting active module containing ACTC1 (encoding the Alpha Actin Cardiac Muscle 1)(Figure 3b). ACTC1 is mainly expressed in developing skeletal muscle, but its expression is also reactivated in diseased mature skeletal muscle, possibly as a sign of regeneration. In this module, we also noticed the presence of the over-expressed gene ABCC8, encoding a modulator of ATP-sensitive potassium channel and insulin release, and involved in the control of contractility and protection of the tissue against calcium overload and fiber damage. An intriguing observation is also the presence of OGDHL, encoding the 2-oxoglutarate deshydrogenase complex component E1-like, which localizes to mitochondria and degrades glucose and glutamate. OGDHL is significantly down-regulated in diseased biopsies. Overall, this active module links a potential reduced mitochondrial activity, frequently described for FSHD1, to decreased response to electrical stimuli (ABCC8) and possible dysfunction of the contractile apparatus associated with over-expression of ACTC1, which might reveal an increased muscle regeneration or immaturity of the contractile apparatus.

In Banerji et al. 2017, the authors highlighted the repression of PAX7 target genes as a hallmark of FSHD1 skeletal muscle [33]. This signature, associated with the activation of the hypoxia pathway, was considered as more robust than the DUX4 signature, which remains variable between studies [42, 32, 44]. We identified 23 active modules with MOGAMUN using this expression dataset (Supplementary Figure S13). The nodes belonging to these modules, as observed for instance in Figure 3c, reveal MAPK-dependent decrease in cell signaling pathways, response to oxydative stress and reduced cell proliferation, as often reported for FSHD1 cells in culture.

We finally applied MOGAMUN to RNA-seq data from myotubes derived from immortalized myoblasts [34]. This RNA-seq study was designed to consider the temporal dimension of gene expression. Genes are classified into 6 categories divided in 3 different groups: up or down regulated in FSHD1; up or down regulated during myogenesis and up or down regulated during FSHD1 myogenesis. One of the main message of this work relates to the suppression of PGC1α (encoded by the PPARGC1A gene) in FSHD1 myotubes as a cause of hypotrophy in FSHD1 myotubes [34]. We applied MOGAMUN only to RNA-seq data obtained from the last time point of the myoblast to myotubes differentiation kinetics (i.e. fully differentiated post-mitotic myotubes) (Material and Method). We identified 17 active modules (Supplementary Figure S14). An interesting module revealed, among other, connections between PPARGC1A and CPT1C (Figure 3d). PPARGC1, down-regulated in FSHD1, is involved in regulating the activities of cAMP response element binding protein (CREB) and nuclear respiratory factors (NRFs). CPT1C, upregulated in FSHD, is involved in muscle glucose uptake. We also observed in the module the presence of MAPK10, required for protection against apoptosis. It is to note that MAPK10 is also identified in a module from FSHD1 immortalized myoblasts (Figure 3c), overall highlighting the existence of connections between specific signalling pathways and chromatin-associated factors previously identified as implicated in the disease (YY1, EP300, CREBBP) [44, 45].

## 4 Discussion

We here designed, compared and applied MOGAMUN, a multi-objective genetic algorithm that is able to detect active modules in multiplex networks. Multiplex biological networks are composed of different layers of physical and functional interactions; each layer has its own meaning, topology and noise. The protein-protein interaction layer, for example, is sparse, but composed of physical binary interactions extracted from curated databases. On the other hand, the co-expression network is very dense, but prone to indirect and spurious interactions. However, altogether, the different layers can provide complementary functional information [17, 18].

We compared MOGAMUN to three different methods, representative of the main algorithms dedicated to the identification of active modules: greedy searches (PinnacleZ), simulated annealing (jActiveModule), and mono-objective genetic algorithm (COSINE). As, to our knowledge, no existing method is able to leverage multiplex networks as inputs, we designed a benchmark for comparison that is based on single networks. In order to have a fair comparison between MOGAMUN and the three other methods, we used the parameters recommended by the authors, except if there were other values that matched our particular selection of parameters (Table 1). In particular, we set the number of subnetworks to be retrieved by jActiveModules to one because there was a single active module. In addition, we set the maximum size per subnetwork to 50 in PinnacleZ. Finally, we set the lambda parameter to 0.5, to have a tradeoff between the weights of the nodes and edges, in COSINE. We demonstrated the performance of MOGAMUN in retrieving modules both densely connected and containing top-scoring nodes, i.e., nodes associated to a high deregulation. However, MOGAMUN running time is, similarly to the other genetic algorithm COSINE, one order of magnitude slower than jActiveModule and PinnacleZ in its current implementation. This running time could be improved by implementing the most computationally demanding tasks (e.g., cross-over) in lower programming languages, like C or Python, or using surrogate-assisted multi-objective evolutionary algorithms.

The extensive analysis of the different FSHD1 datasets highlighted the reduced proliferation and increased apoptosis of cells from FSHD1 patients and led to the identification of novel genes in this different pathways by linking cell defects to factors involved in muscle function. It further revealed consistencies in biological processes identified by different teams in their respective models but also some putative discrepancies in the interpretation of disease-associated biological processes depending on the type of samples used (biopsies of muscle unaffected in the disease, immortalized or transduced proliferative myoblasts or post-mitotic myotubes). Overall, this also reveals that MOGAMUN can be applied to identify disease-associated biological processes in rare diseases for which the number of samples is limited, and also to compare the processes identified in different datasets.

We applied here MOGAMUN to identify active module from the integration of RNA-seq expression data into multiplex networks. However, it is to note that any type of molecular profile associated to *p*-values can be integrated on the networks, such as *p*-values obtained from a GWAS, from phenotypic hit screening, or from proteomics profiling.

## ACKNOWLEDGMENTS

E.M.N.T was supported by CONACYT PhD fellowship. The project leading to this publication has received funding from the Excellence Initiative of Aix-Marseille University - A*Midex, a French “Investissements d’Avenir” programme. We thank Elisabeth Remy (Aix Marseille Univ, CNRS, I2M, Marseille Institute of Mathematics, Marseille, France), and Claude Pasquier (Univ. Nice Sophia Antipolis, CNRS, I3S, Nice, France), for useful discussions.

## Notes

### Competing Interest Statement

The authors have declared no competing interest.

## References

[1] J. Reimand, R. Isserlin, V. Voisin, M. Kucera, C. Tannus-Lopes, A. Rostamianfar, …, and D. Merico. Pathway enrichment analysis and visualization of omics data using g: Profiler, gsea, cytoscape and enrichmentmap. Nature protocols 14, 2019.

[2] K. Mitra, A. R. Carvunis, S. K. Ramesh, and T. Ideker. Integrative approaches for finding modular structure in biological networks. Nature Reviews Genetics 14, 2013.

[3] T. Ideker, O. Ozier, B. Schwikowski, and A. F. Siegel. Discovering regulatory and signalling circuits in molecular interaction networks. Bioinformatics 18, 2002.

[4] D. Li, Z. Pan, G. Hu, Z. Zhu, and S. He. Active module identification in intracellular networks using a memetic algorithm with a new binary decoding scheme. BMC genomics 18, 2017.

[5] W. Chen, J. Liu, and S. He. Prior knowledge guided active modules identification: an integrated multi-objective approach. BMC systems biology 11, 2017.

[6] D. Li, J. B. Brown, L. Orsini, Z. Pan, G. Hu, and S. He. Moda: Module differential analysis for weighted gene co-expression network. arXiv preprint arXiv:1605.04739, 2016.

[7] B. Zhang and S. Horvath. A general framework for weighted gene co-expression network analysis. Statistical applications in genetics and molecular biology 4, 2005.

[8] K. Kusonmano, M. K. Halle, E. Wik, E. A. Hoivik, C. Krakstad, K. K. Mauland, …, and A. M. Oyan. Identification of highly connected and differentially expressed gene subnetworks in metastasizing endometrial cancer. Plos one 13, 2018.

[9] H. Nguyen, S. Shrestha, D. Tran, A. Shafi, S. Draghici, and T. Nguyen. A comprehensive survey of tools and software for active subnetwork identification. Frontiers in genetics 10, 2019.

[10] H. Y. Chuang, E. Lee, Y. T. Liu, D. Lee, and T. Ideker. Network-based classification of breast cancer metastasis. Molecular systems biology 3, 2007.

[11] I. Ulitsky and R. Shamir. Identification of functional modules using network topology and high-throughput data. BMC systems biology 1, 2007.

[12] H. Ma, E. E. Schadt, L. M. Kaplan, and H. Zhao. Cosine: Condition-specific subnetwork identification using a global optimization method. Bioinformatics 27, 2011.

[13] D. Muraro and A. Simmons. An integrative analysis of gene expression and molecular interaction data to identify dys-regulated subnetworks in inflammatory bowel disease. BMC bioinformatics 17, 2016.

[14] O. Ozisik, B. Bakir-Gungor, B. Diri, and O. Ugur Sezerman. Active subnetwork ga: a two stage genetic algorithm approach to active subnetwork search. A Current Bioinformatics 12, 2017.

[15] Y. Liu, M. Brossard, D. Roqueiro, P. Margaritte-Jeannin, C. Sarnowski, E. Bouzigon, and F. Demenais. Sigmod: an exact and efficient method to identify a strongly interconnected disease-associated module in a gene network. Bioinformatics 33, 2017.

[16] F. Battiston, V. Nicosia, and V. Latora. Structural measures for multiplex networks. Physical Review E 89, 2014.

[17] A. Valdeolivas, L. Tichit, C. Navarro, S. Perrin, G. Odelin, N. Levy, …, and A. Baudot. Random walk with restart on multiplex and heterogeneous biological networks. Bioinformatics 35, 2018.

[18] G. Didier, C. Brun, and A. Baudot. Identifying communities from multiplex biological networks. PeerJ 3, 2015.

[19] A. Halu, M. De Domenico, A. Arenas, and A. Sharma. The multiplex network of human diseases. NPJ systems biology and applications 5, 2019.

[20] L. Bennett, A. Kittas, G. Muirhead, L. G. Papageorgiou, and S. Tsoka. Detection of composite communities in multiplex biological networks. Scientific reports 5, 2015.

[21] G. Mangioni, G. Jurman, and M. De-Domenico. Multilayer flows in molecular networks identify biological modules in the human proteome. IEEE Transactions on Network Science and Engineering 7, 2020.

[22] R. Kanawati. Multiplex network mining: A brief survey. IEEE Intelligent Informatics Bulletin 16, 2015.

[23] K. Deb, S. Agrawal, A. Pratap, and T. Meyarivan. A fast elitist non-dominated sorting genetic algorithm for multi-objective optimization: Nsga-ii. International conference on parallel problem solving from nature, 2000.

[24] K. Deb. Multi-objective optimization. In Search methodologies., chapter 15, pages 403–449. Springer, Boston, MA., 2014.

[25] T. Blickle. Tournament selection. Evolutionary computation 1, 2000.

[26] S. Choobdar, M. E. Ahsen, J. Crawford, M. Tomasoni, T. Fang, D. Lamparter, …, and T. Natoli. Assessment of network module identification across complex diseases. Nature methods 16, 2019.

[27] R. Batra, N. Alcaraz, K. Gitzhofer, J. Pauling, H. J. Ditzel, M. Hellmuth, and M. List. On the performance of de novo pathway enrichment. NPJ systems biology and applications 3, 2017.

[28] T. S. Keshava Prasad, R. Goel, K. Kandasamy, S. Keerthikumar, S. Kumar, S. Mathivanan, …, and L. Balakrishnan. Human protein reference database—2009 update. Nucleic acids research 37, 2008.

[29] N. del Toro, M. Dumousseau, S. Orchard, R. C. Jimenez, E. Galeota, G. Launay, …, and H. Hermjakob. A new reference implementation of the psicquic web service. Nucleic acids research 41, 2013.

[30] T. Rolland, M. Tasan, B. Charloteaux, S. J. Pevzner, Q. Zhong, N. Sahni, …, and A. Kamburov. A proteome-scale map of the human interactome network. Cell 159, 2014.

[31] M. D. Robinson, D. J. McCarthy, and G. K. Smyth. edger: a bioconductor package for differential expression analysis of digital gene expression data. Bioinformatics 26, 2010.

[32] Z. Yao, L. Snider, J. Balog, R. J. Lemmers, S. M. Van Der Maarel, R. Tawil, and S. J. Tapscott. Dux4-induced gene expression is the major molecular signature in fshd skeletal muscle. Human molecular genetics 23, 2014.

[33] C. R. Banerji, M. Panamarova, H. Hebaishi, R. B. White, F. Relaix, S. Severini, and P. S. Zammit. Pax7 target genes are globally repressed in facioscapulohumeral muscular dystrophy skeletal muscle. Nature communications 8, 2017.

[34] C. R. Banerji, M. Panamarova, J. Pruller, N. Figeac, H. Hebaishi, E. Fidanis, …, and P. S. Zammit. Dynamic transcriptomic analysis reveals suppression of pgc1 *α*/err *α* drives perturbed myogenesis in facioscapulohumeral muscular dystrophy. Human molecular genetics 28, 2019.

[35] R. Edgar, M. Domrachev, and A. E. Lash. Gene expression omnibus: Ncbi gene expression and hybridization array data repository. Nucleic acids research 30, 2002.

[36] J. M. Young, J. L. Whiddon, Z. Yao, B. Kasinathan, L. Snider, L. N. Geng, …, and S. J. Tapscott. Dux4 binding to retroelements creates promoters that are active in fshd muscle and testis. PLoS genetics 9, 2013.

[37] Y. D. Krom, J. Dumonceaux, K. Mamchaoui, B. den Hamer, V. Mariot, E. Negroni, …, and B. G. van Engelen. Generation of isogenic d4z4 contracted and noncontracted immortal muscle cell clones from a mosaic patient: a cellular model for fshd. The American journal of pathology 181, 2012.

[38] S. Homma, J. C. Chen, F. Rahimov, M. L. Beermann, K. Hanger, G. M. Bibat, …, and J. B. Miller. A unique library of myogenic cells from facioscapulohumeral muscular dystrophy subjects and unaffected relatives: family, disease and cell function. European journal of human genetics 20, 2012.

[39] G. Sales, E. Calura, D. Cavalieri, and C. Romualdi. graphite -a bioconductor package to convert pathway topology to gene network. BMC bioinformatics 13, 2012.

[40] M. Uhlén, L. Fagerberg, B. M. Hallström, C. Lindskog, P. Oksvold, A. Mardinoglu, …, and I. Olsson. Tissue-based map of the human proteome. Science 347, 2015.

[41] K. Nguyen, N. Broucqsault, C. Chaix, S. Roche, J. D. Robin, C. Vovan, …, and C. Barnérias. Deciphering the complexity of the 4q and 10q subtelomeres by molecular combing in healthy individuals and patients with facioscapulohumeral dystrophy. Journal of medical genetics 56, 2019.

[42] L. N. Geng, Z. Yao, L. Snider, A. P. Fong, J. N. Cech, J. M. Young, …, and S. J. Tapscott. Dux4 activates germline genes, retroelements, and immune mediators: implications for facioscapulohumeral dystrophy. Developmental cell 22, 2012.

[43] A. Pakula, J. Schneider, J. Janke, U. Zacharias, H. Schulz, N. Hübner,.., and M. Boschmann. Altered expression of cyclin a 1 in muscle of patients with facioscapulohumeral muscle dystrophy (fshd-1). PloS one 8, 2013.

[44] S. H. Choi, M. D. Gearhart, Z. Cui, D. Bosnakovski, M. Kim, N. Schennum, and M. Kyba. Dux4 recruits p300/cbp through its c-terminus and induces global h3k27 acetylation changes. Nucleic acids research 44, 2016.

[45] D. Gabellini, M. R. Green, and R. Tupler. Inappropriate gene activation in fshd: a repressor complex binds a chromosomal repeat deleted in dystrophic muscle. Cell 110, 2002.

