## Supplementary data for "A Multi-Objective Genetic Algorithm to Find Active Modules in Multiplex Biological Networks"

### A Multi-Objective Genetic Algorithm to Find Active Modules from Multiplex Biological Networks: Supplementary material


#### Contents


**Figure S1** **2**

**Figure S2** **2**

**Figure S3** **3**

**Figure S4** **3**

**Figure S5** **4**

**Figure S6** **4**

**Figure S7** **5**

**Figure S8** **5**

**Figure S9** **6**

**Figure S10** **6**

**Figure S11** **14**

**Figure S12** **18**

**Figure S13** **30**

**Figure S14** **38**

**Table S1** **48**

**Table S2** **48**

**Table S3** **49**

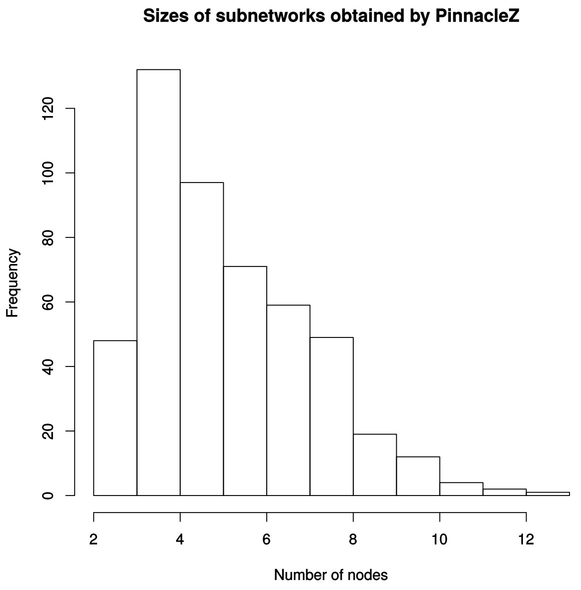


Figure S1: Sizes of the subnetworks identified by PinnacleZ in the experiment using the network $PPI\_1$ and the simulated data with normal distribution

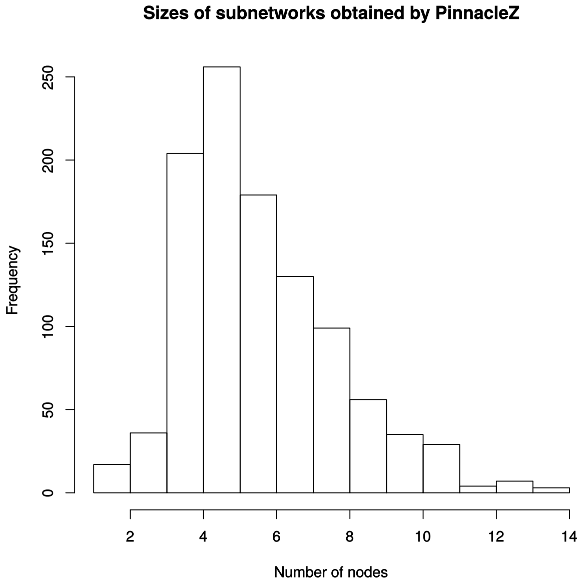


Figure S2: Sizes of the subnetworks identified by PinnacleZ in the experiment using the network $PPI\_2$ and the sampled data from RNA-Seq TCGA breast cancer dataset

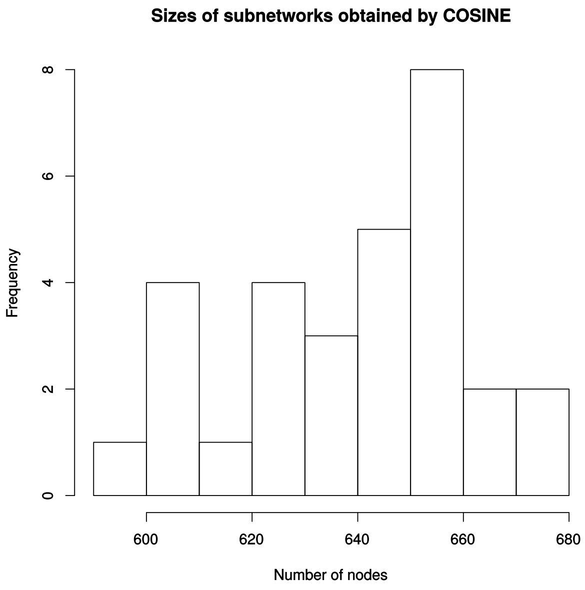


Figure S3: Sizes of the subnetworks identified by COSINE in the experiment using the network $PPI\_1$ and the simulated data with normal distribution

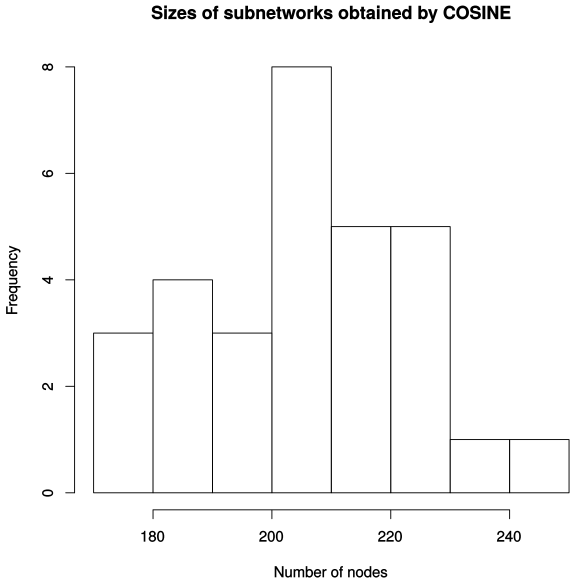


Figure S4: Sizes of the subnetworks identified by COSINE in the experiment using the network $PPI\_2$ and the sampled data from RNA-Seq TCGA breast cancer dataset

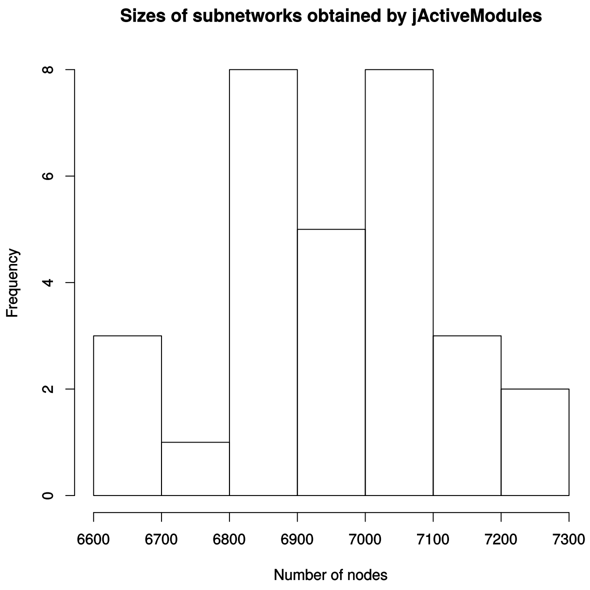


Figure S5: Sizes of the subnetworks identified by jActiveModules in the experiment using the network $PPI\_1$ and the simulated data with normal distribution

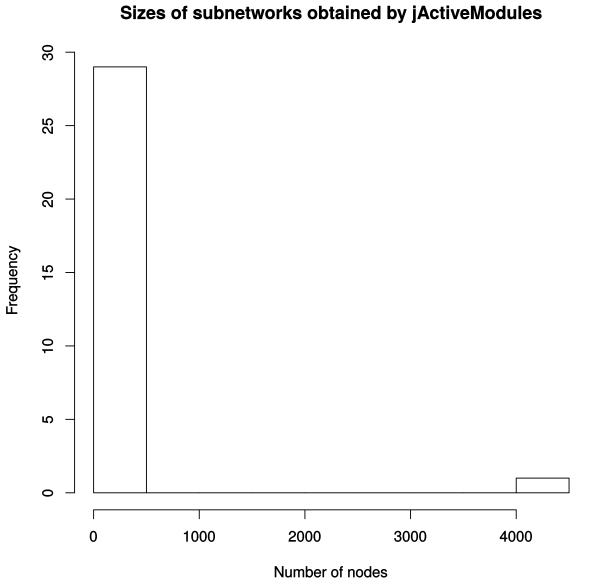


Figure S6: Sizes of the subnetworks identified by jActiveModules in the experiment using the network $PPI\_2$ and the sampled data from RNA-Seq TCGA breast cancer dataset

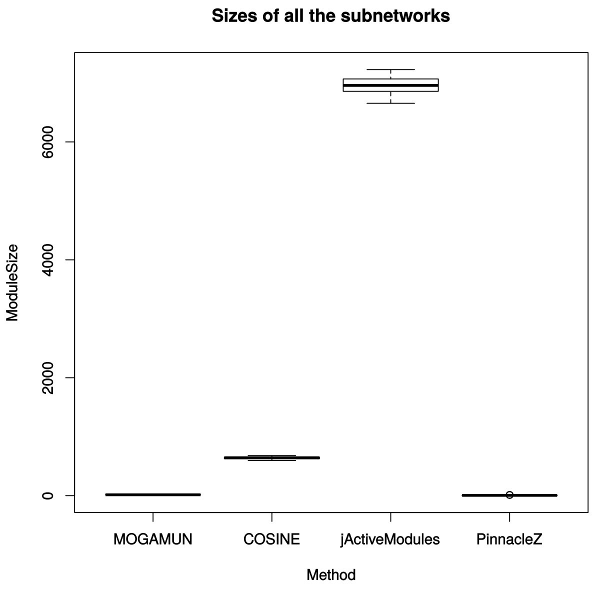


Figure S7: Sizes of the subnetworks identified by all the methods in the experiment using the network $PPI\_1$ and the simulated data with normal distribution

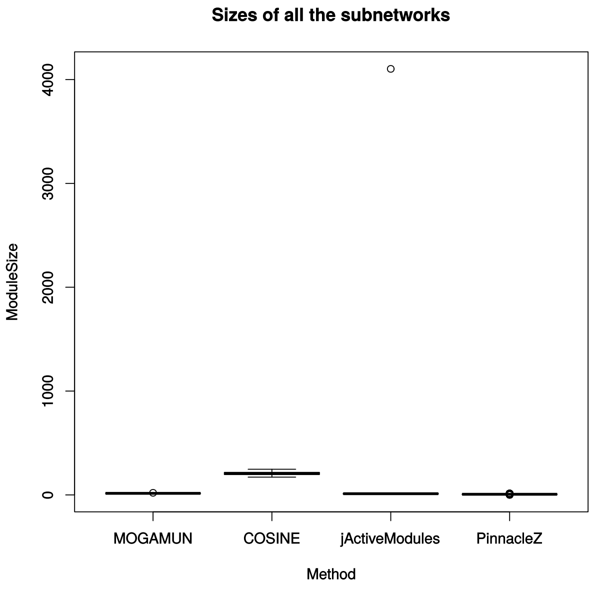


Figure S8: Sizes of the subnetworks identified by all the methods in the experiment using the network $PPI\_2$ and the sampled data from RNA-Seq TCGA breast cancer dataset

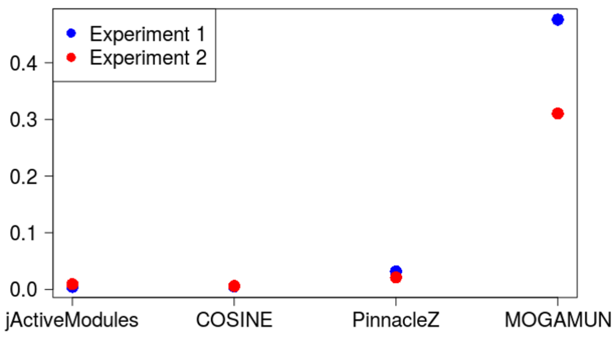


Figure S9: $F_1$ score values of jActiveModules, COSINE, PinnacleZ, and MOGAMUN corresponding to the two experiments from the benchmark

### Figure S10

Eighteen active modules obtained by applying MOGAMUN on Yao's biopsies dataset [1] (see Table S1 for the list of samples). The color of the nodes represents the fold-change, where green and red nodes correspond to under- and over-expressed genes, respectively. Nodes with bold black border correspond to genes significantly differentially expressed ($FDR < 0.05$ and absolute $log_2$ fold-change $> 1$). Blue and white nodes correspond to genes with no associated transcriptomics data and no deregulation, respectively.

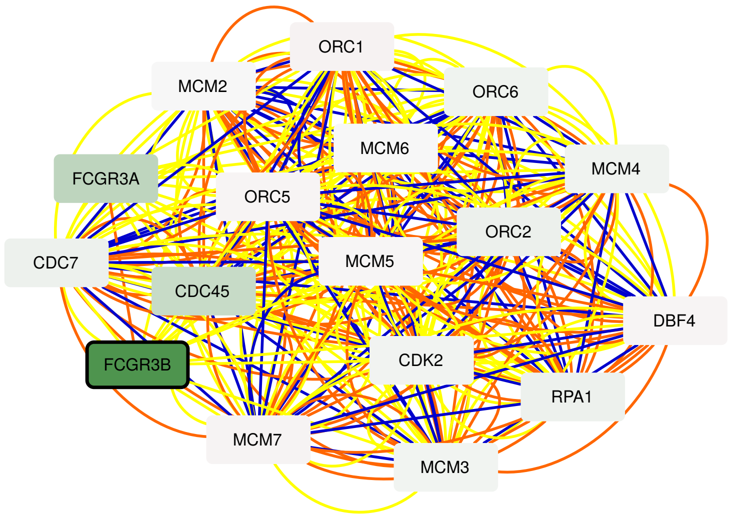


Figure S10.1. Yao's dataset, biopsies: Active module 1

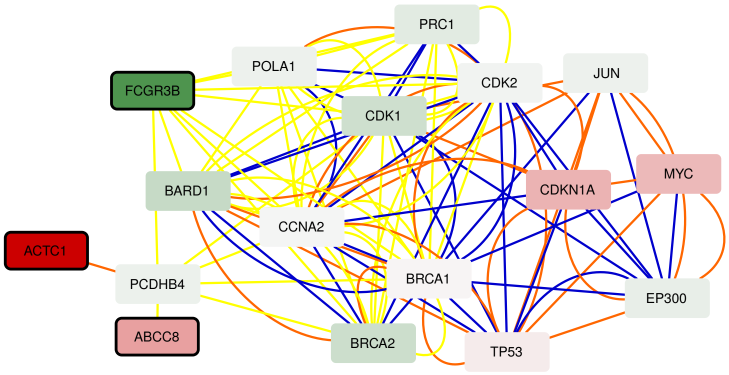


Figure S10.2. Yao's dataset, biopsies: Active module 2

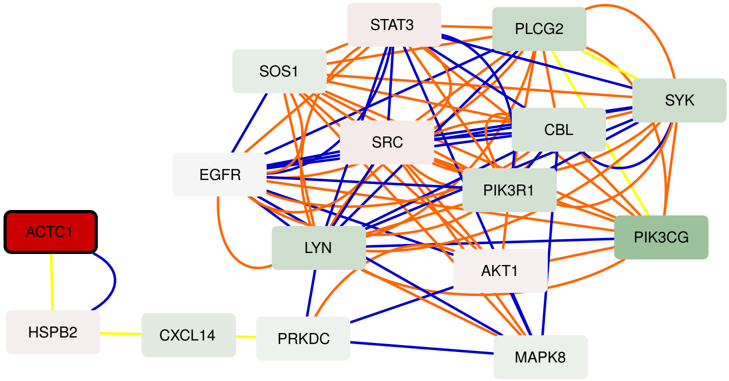


Figure S10.3. Yao's dataset, biopsies: Active module 3

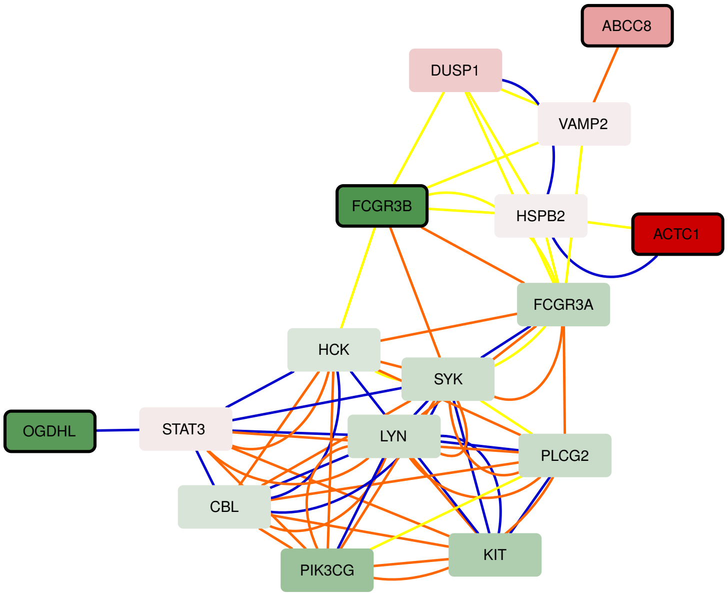


Figure S10.4. Yao's dataset, biopsies: Active module 4

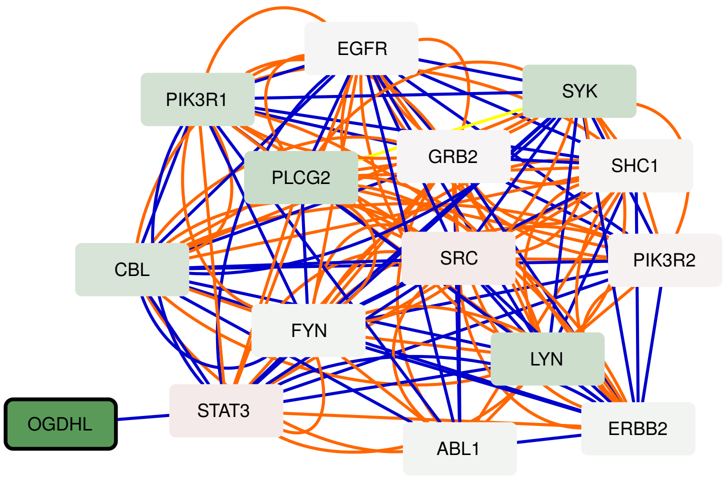


Figure S10.5. Yao's dataset, biopsies: Active module 5

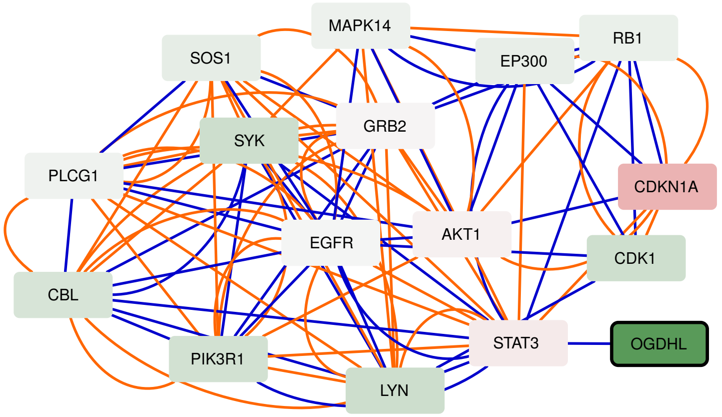


Figure S10.6. Yao's dataset, biopsies: Active module 6

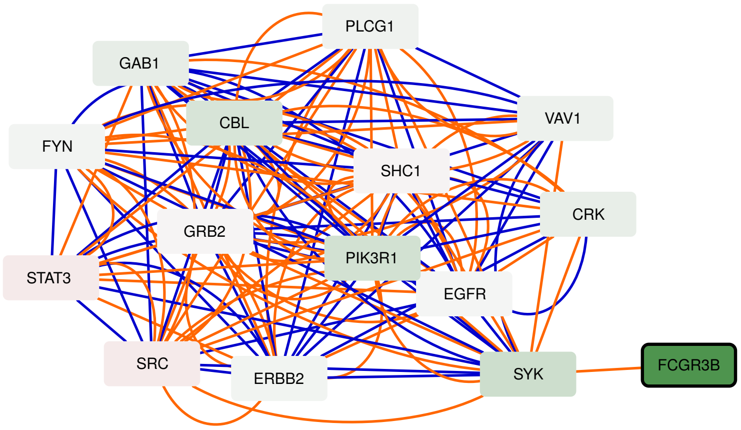


Figure S10.7. Yao's dataset, biopsies: Active module 7

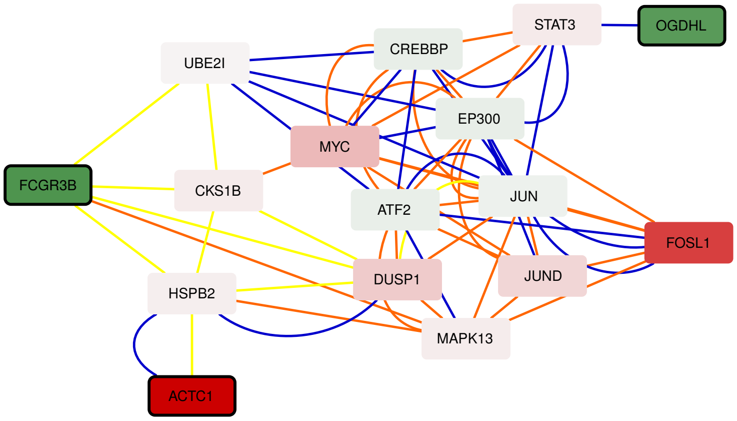


Figure S10.8. Yao's dataset, biopsies: Active module 8

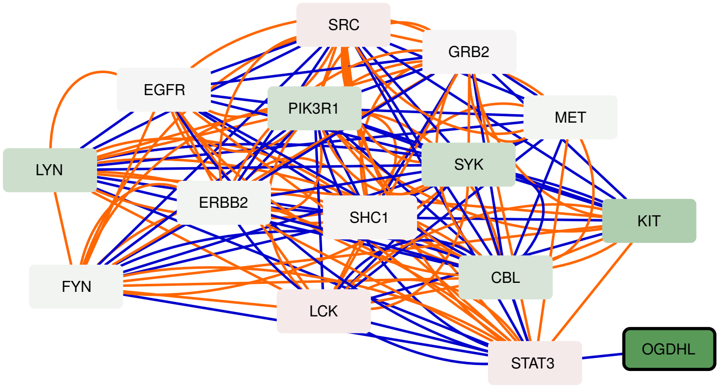


Figure S10.9. Yao's dataset, biopsies: Active module 9

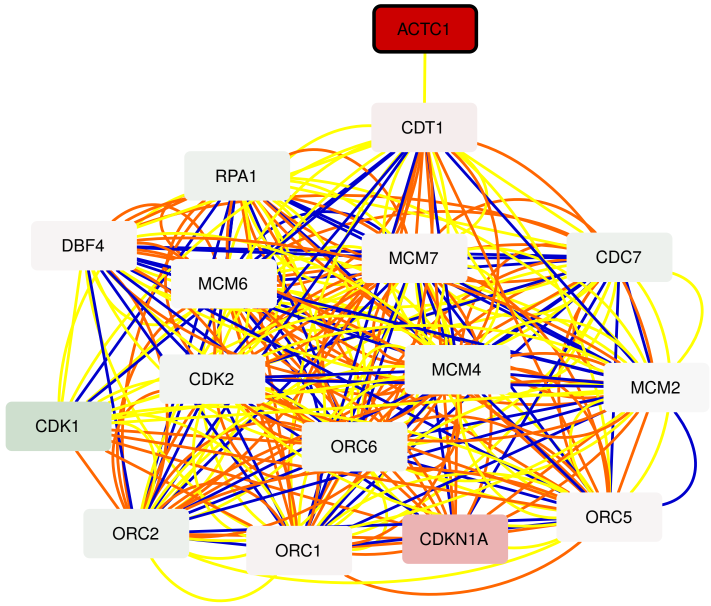


Figure S10.10. Yao's dataset, biopsies: Active module 10

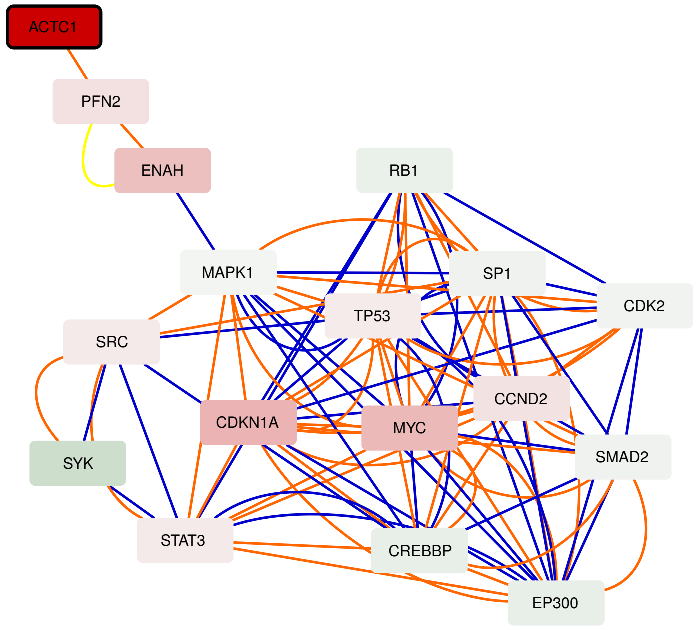


Figure S10.11. Yao's dataset, biopsies: Active module 11

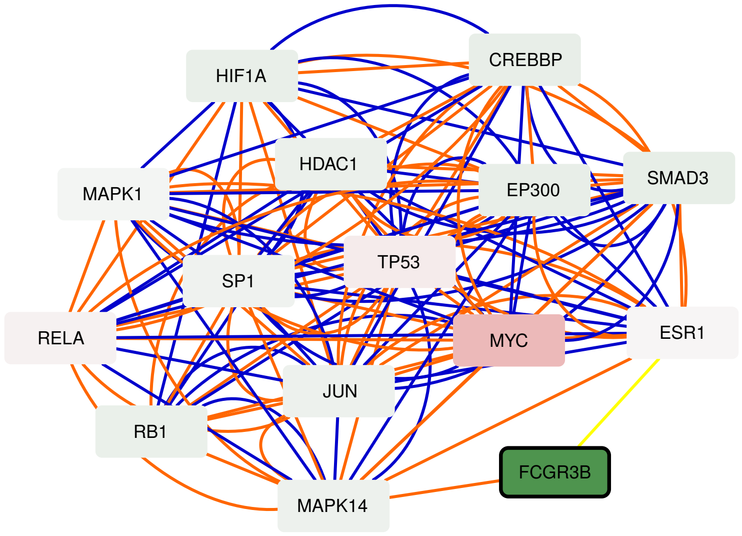


Figure S10.12. Yao's dataset, biopsies: Active module 12

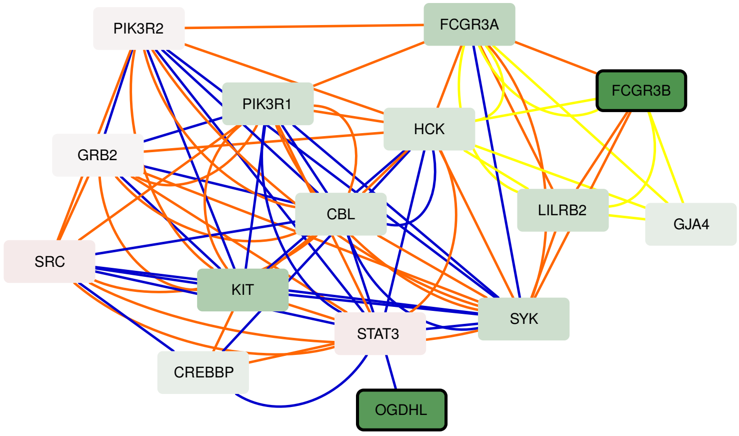


Figure S10.13. Yao's dataset, biopsies: Active module 13

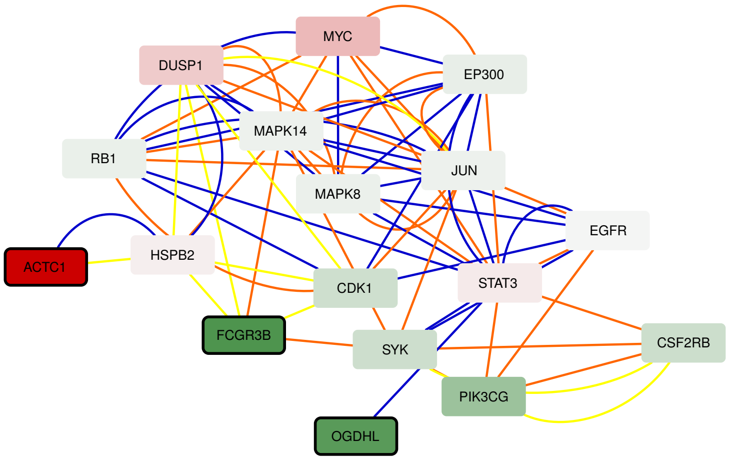


Figure S10.14. Yao's dataset, biopsies: Active module 14

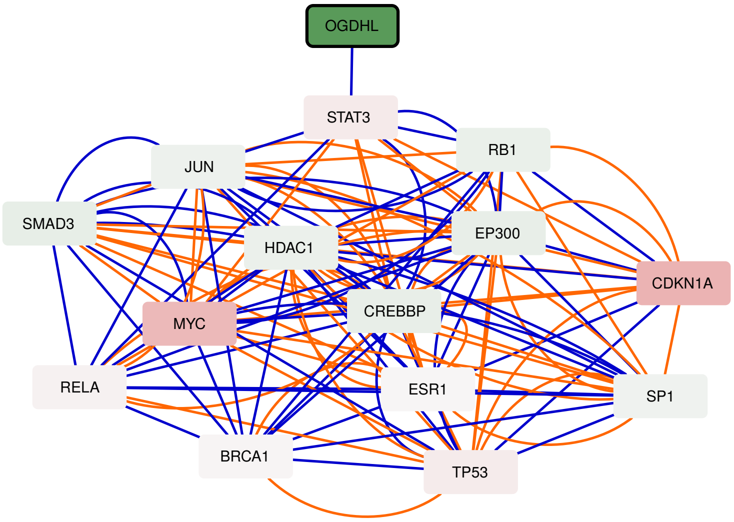


Figure S10.15. Yao's dataset, biopsies: Active module 15

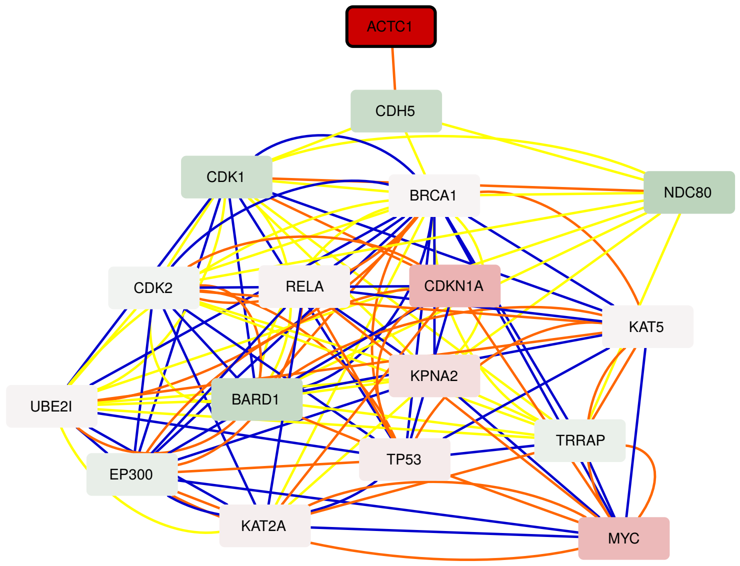


Figure S10.16. Yao's dataset, biopsies: Active module 16

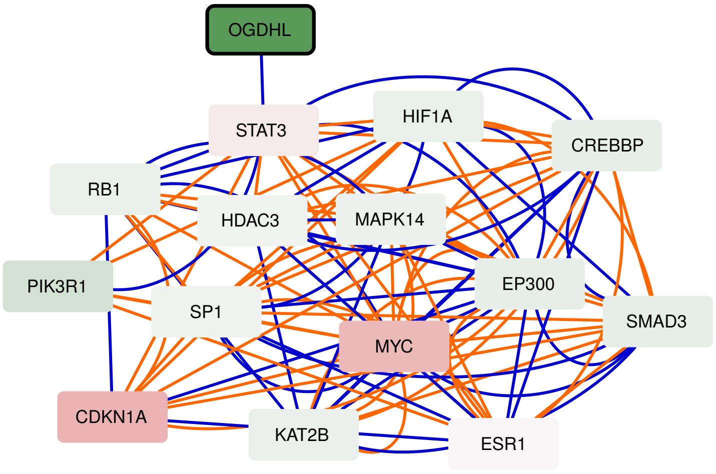


Figure S10.17. Yao's dataset, biopsies: Active module 17

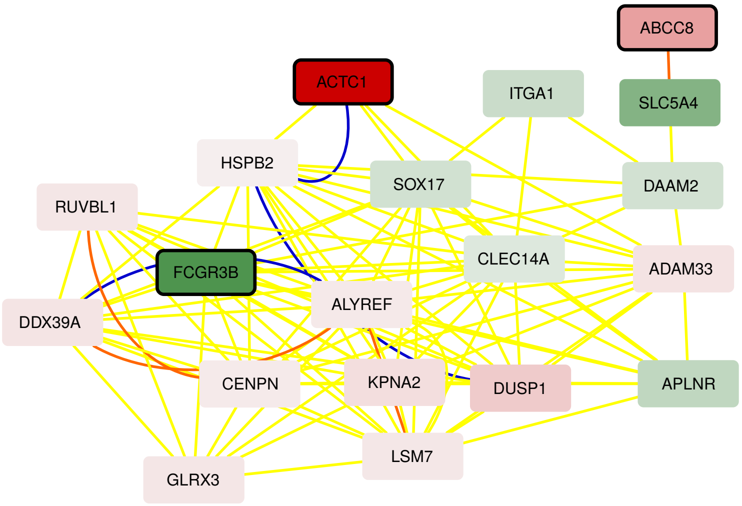


Figure S10.18. Yao's dataset, biopsies: Active module 18

### Figure S11

Ten active modules obtained by applying MOGAMUN on Yao's myoblasts dataset [1] (see Table S1 for the list of samples). The color of the nodes represents the fold-change, where green and red nodes correspond to under- and over-expressed genes, respectively. Nodes with bold black border correspond to genes significantly differentially expressed ($FDR < 0.05$ and absolute $log_2$ fold-change $> 1$). Blue and white nodes correspond to genes with no associated transcriptomics data and no deregulation, respectively.

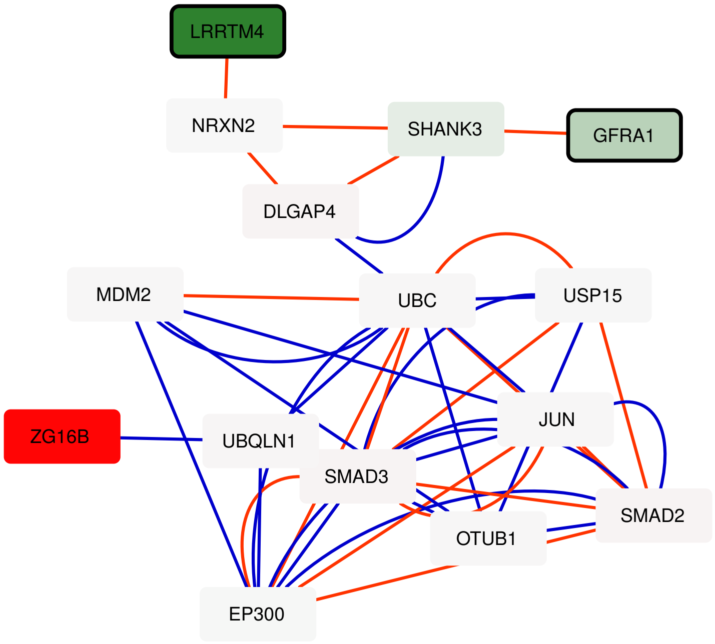


Figure S11.1. Yao's dataset, myoblasts: Active module 1

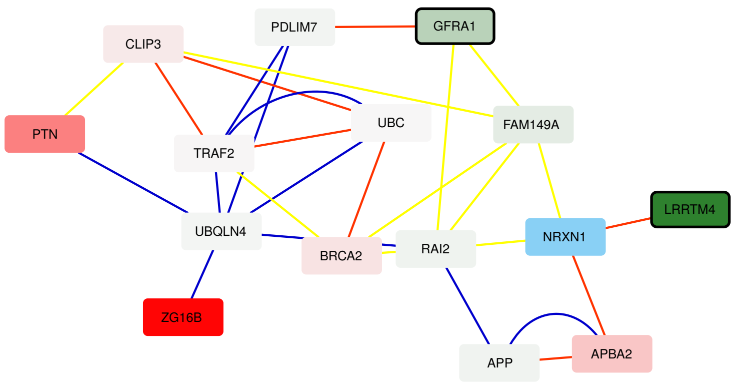


Figure S11.2. Yao's dataset, myoblasts: Active module 2

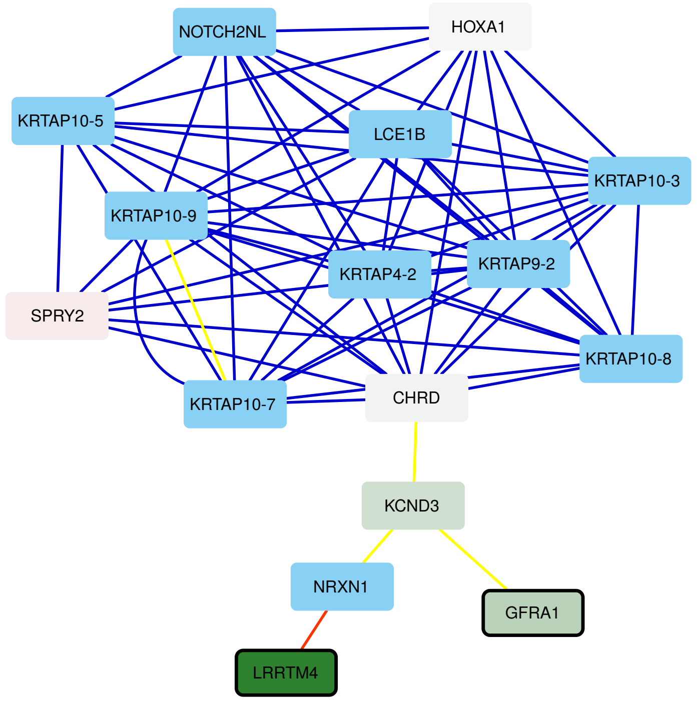


Figure S11.3. Yao's dataset, myoblasts: Active module 3

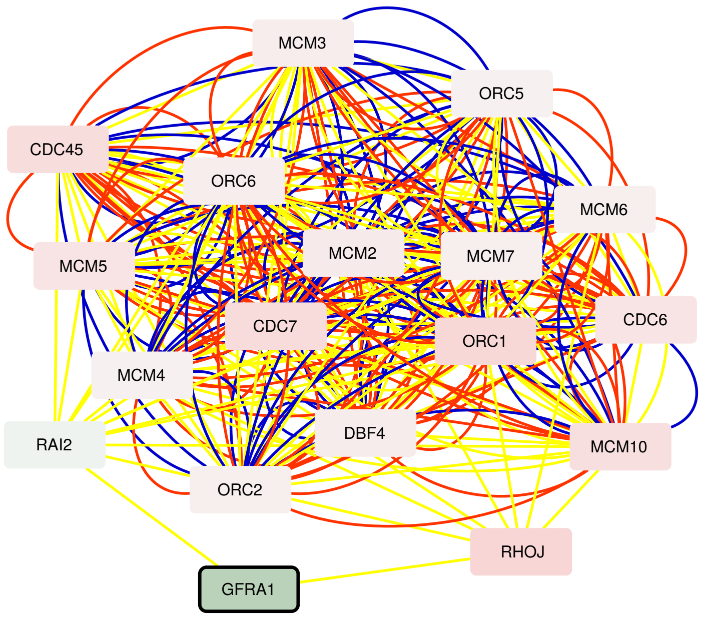


Figure S11.4. Yao's dataset, myoblasts: Active module 4

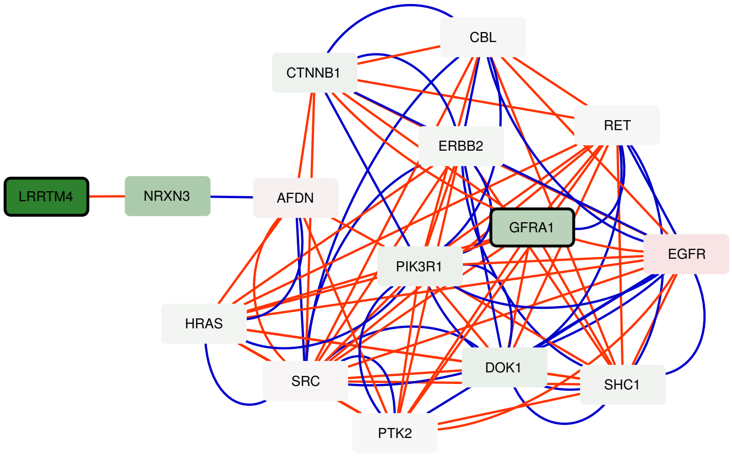


Figure S11.5. Yao's dataset, myoblasts: Active module 5

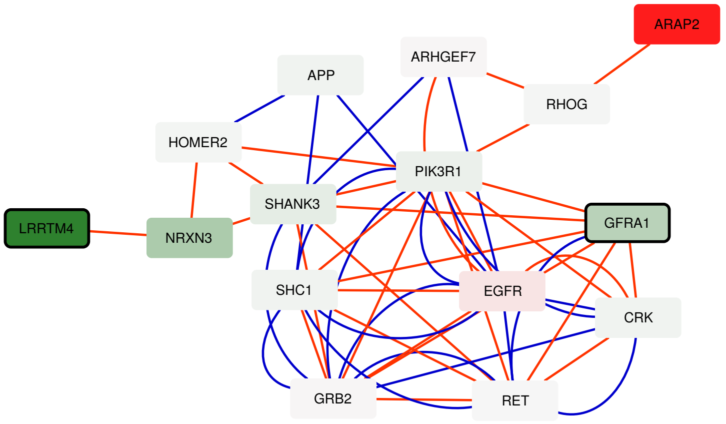


Figure S11.6. Yao's dataset, myoblasts: Active module 6

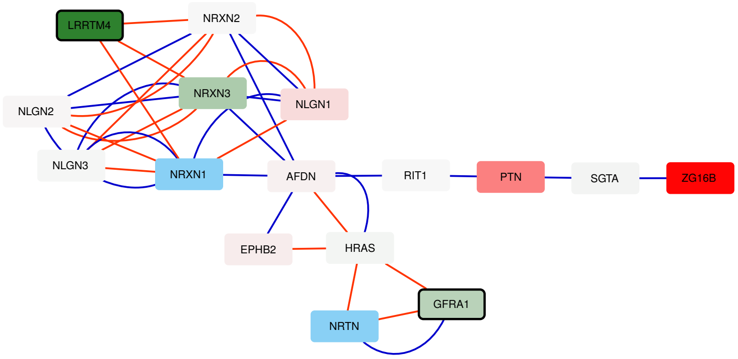


Figure S11.7. Yao's dataset, myoblasts: Active module 7

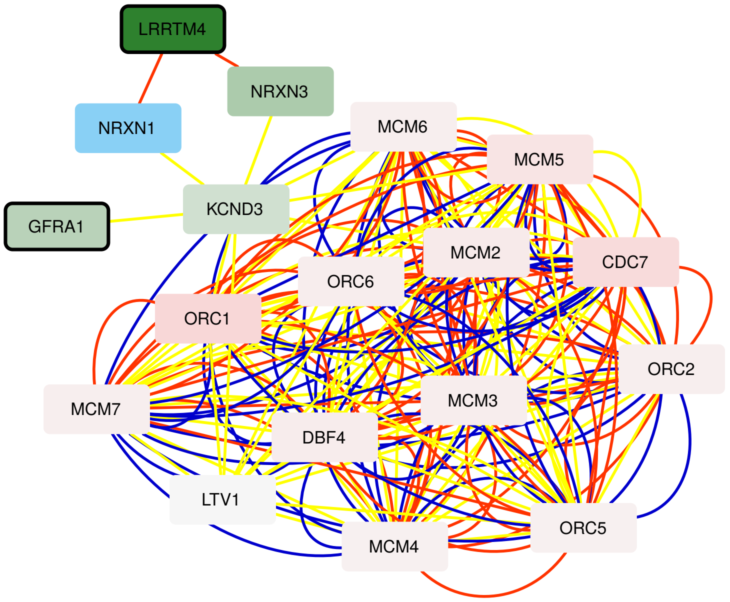


Figure S11.8. Yao's dataset, myoblasts: Active module 8

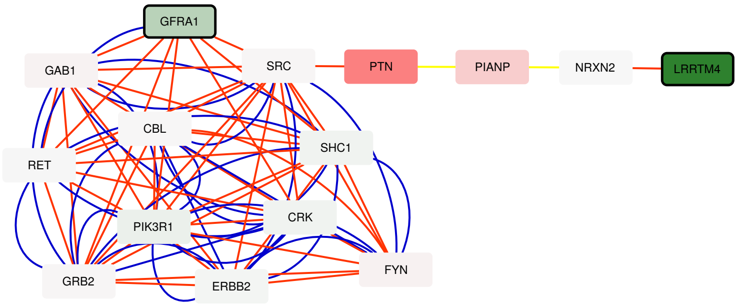


Figure S11.9. Yao's dataset, myoblasts: Active module 9

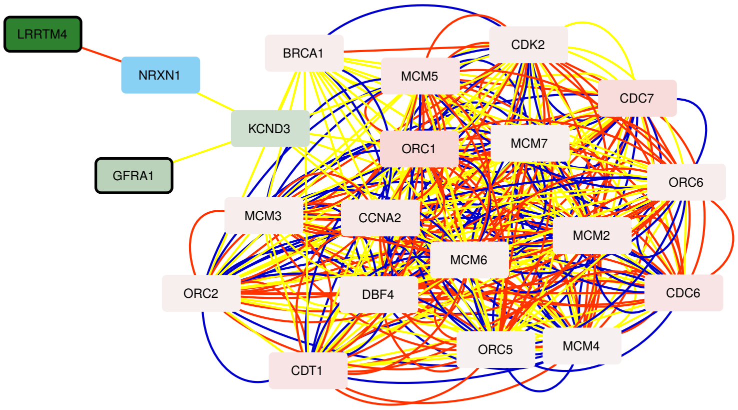


Figure S11.10. Yao's dataset, myoblasts: Active module 10

### Figure S12

Twenty three active modules obtained by applying MOGAMUN on Yao's myotubes dataset [1] (see Table S1 for the list of samples). The color of the nodes represents the fold-change, where green and red nodes correspond to under- and over-expressed genes, respectively. Nodes with bold black border correspond to genes significantly differentially expressed ($FDR < 0.05$ and absolute $log_2$ fold-change $> 1$). Blue and white nodes correspond to genes with no associated transcriptomics data and no deregulation, respectively.

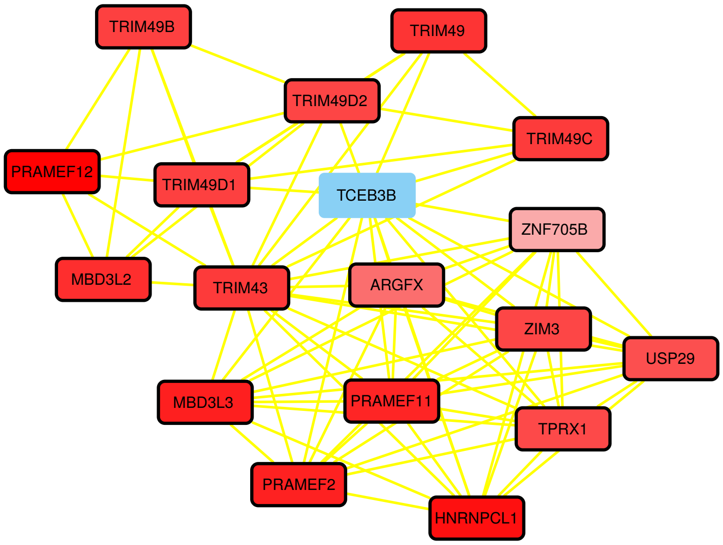


Figure S12.1. Yao's dataset, myotubes: Active module 1

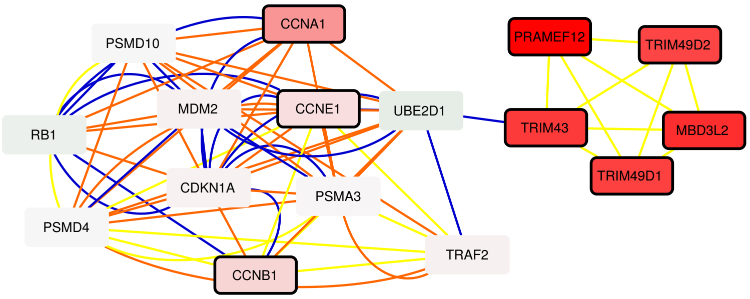


Figure S12.2. Yao's dataset, myotubes: Active module 2

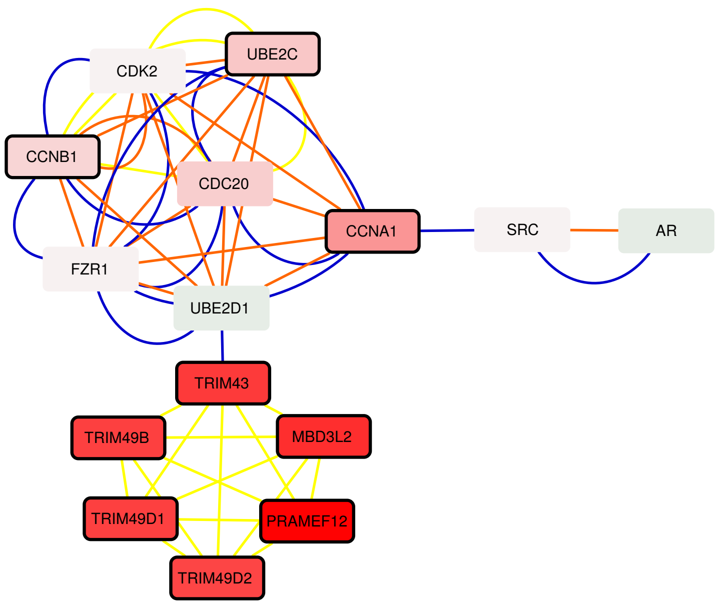


Figure S12.3. Yao's dataset, myotubes: Active module 3

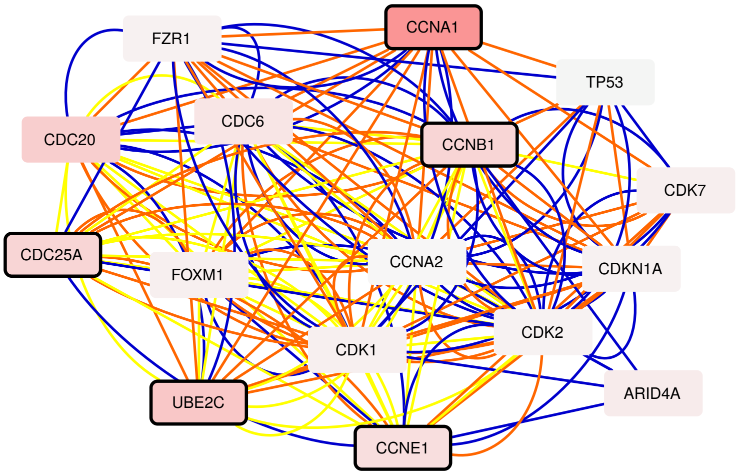


Figure S12.4. Yao's dataset, myotubes: Active module 4

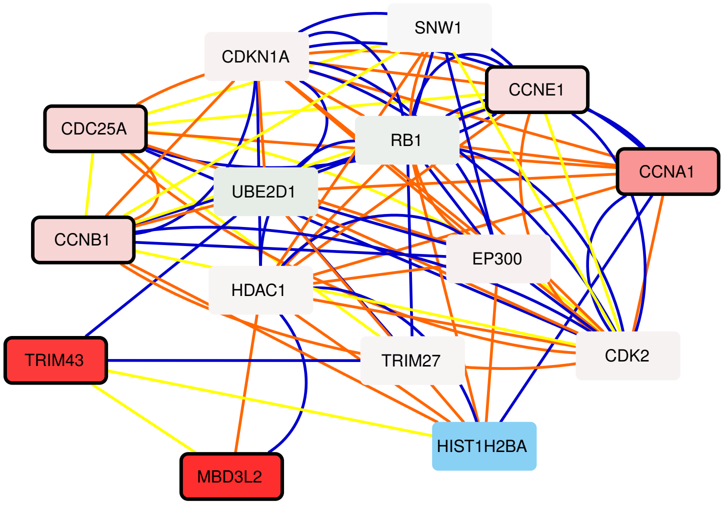


Figure S12.5. Yao's dataset, myotubes: Active module 5

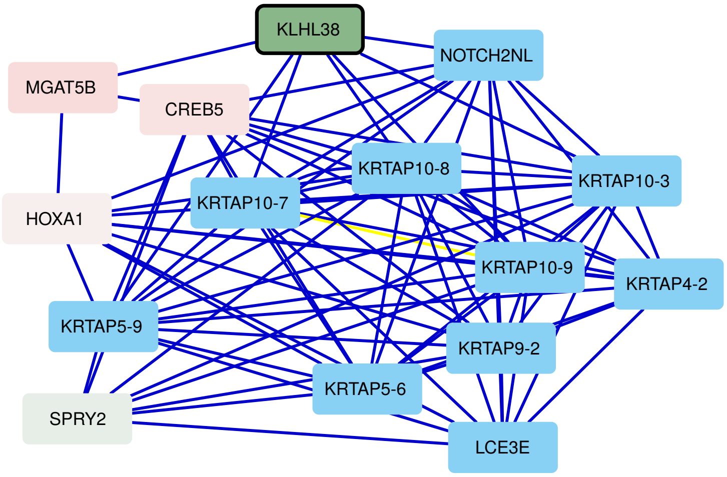


Figure S12.6. Yao's dataset, myotubes: Active module 6

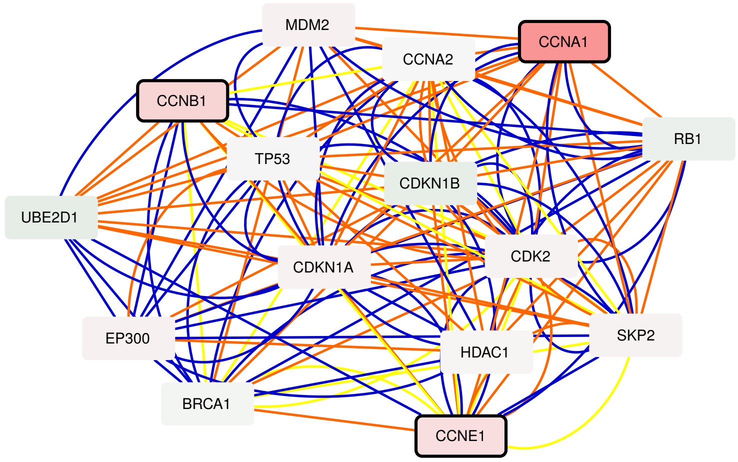


Figure S12.7. Yao's dataset, myotubes: Active module 7

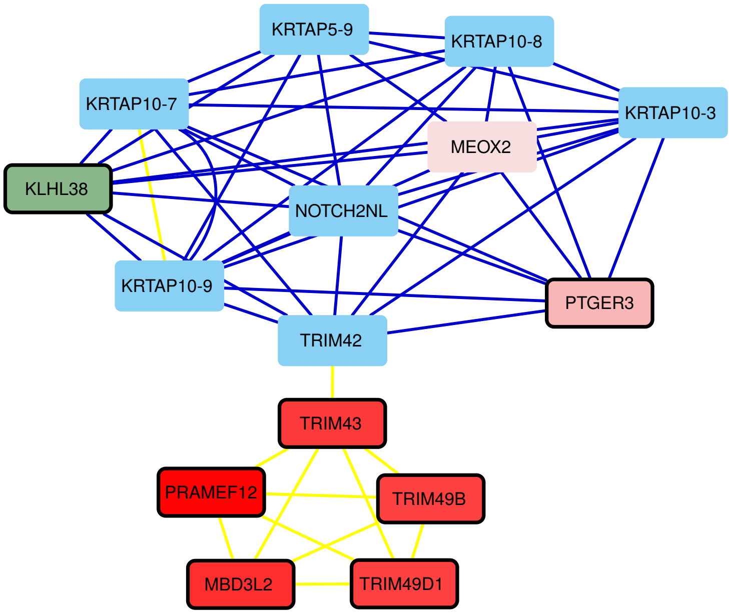


Figure S12.8. Yao's dataset, myotubes: Active module 8

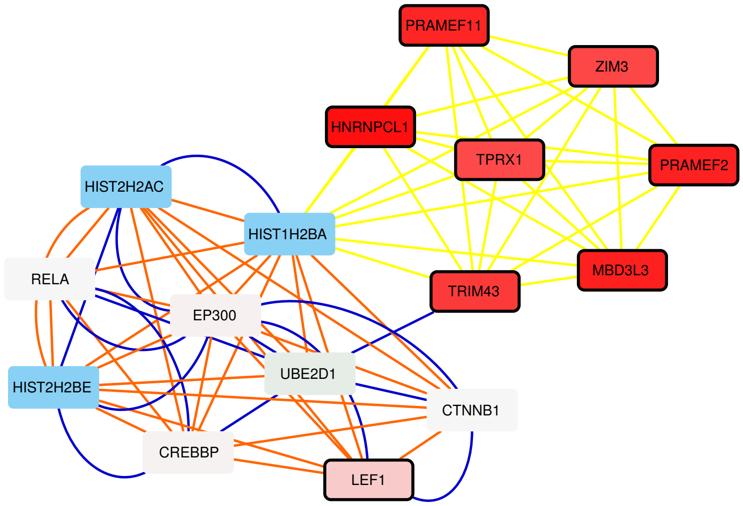


Figure S12.9. Yao's dataset, myotubes: Active module 9

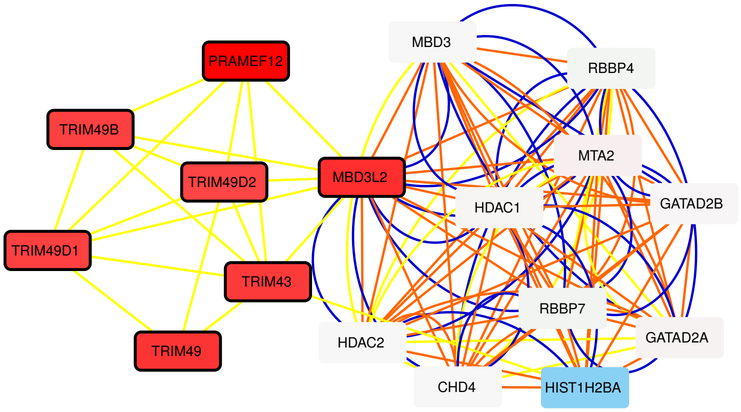


Figure S12.10. Yao's dataset, myotubes: Active module 10

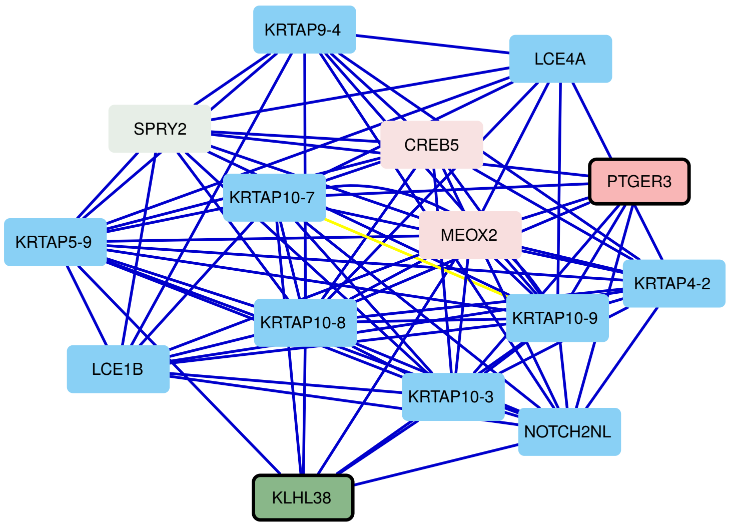


Figure S12.11. Yao's dataset, myotubes: Active module 11

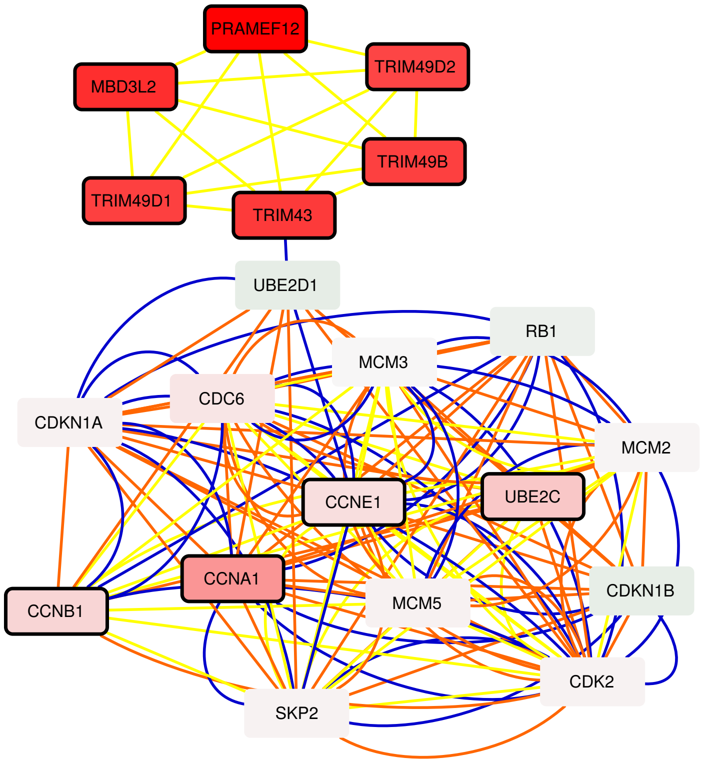


Figure S12.12. Yao's dataset, myotubes: Active module 12

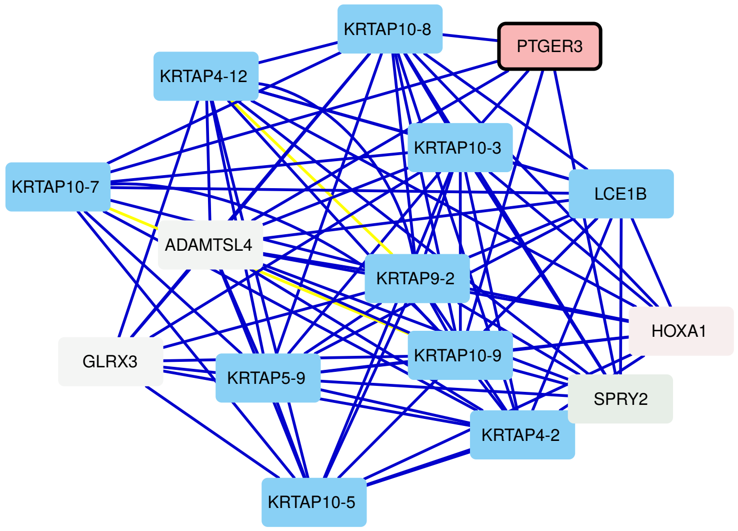


Figure S12.13. Yao's dataset, myotubes: Active module 13

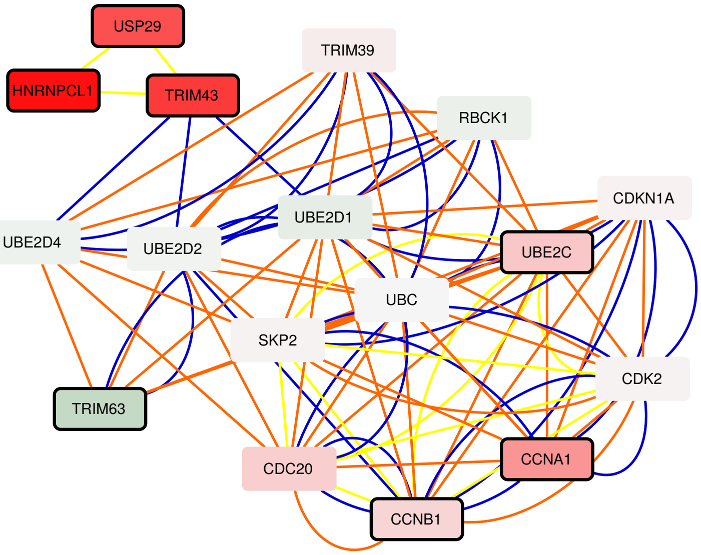


Figure S12.14. Yao's dataset, myotubes: Active module 14

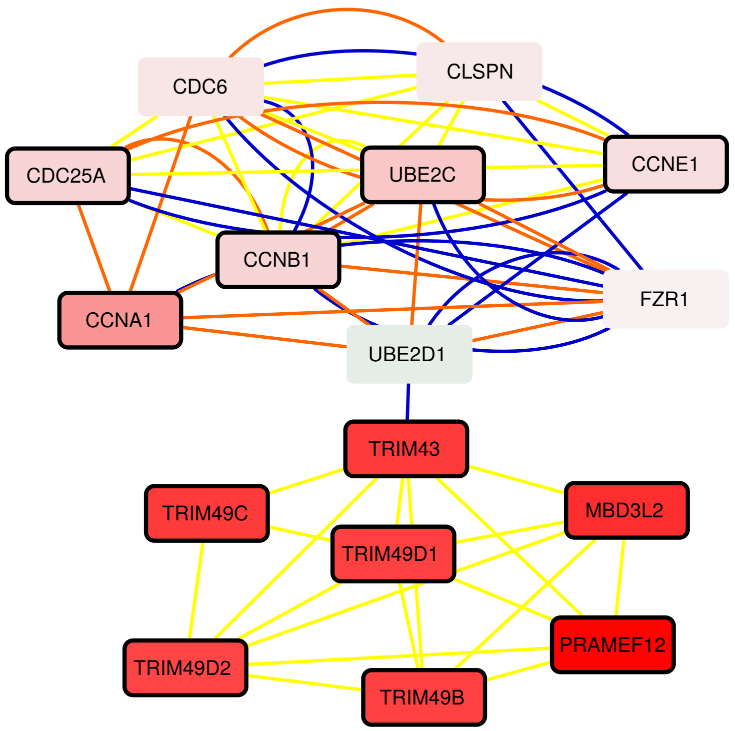


Figure S12.15. Yao's dataset, myotubes: Active module 15

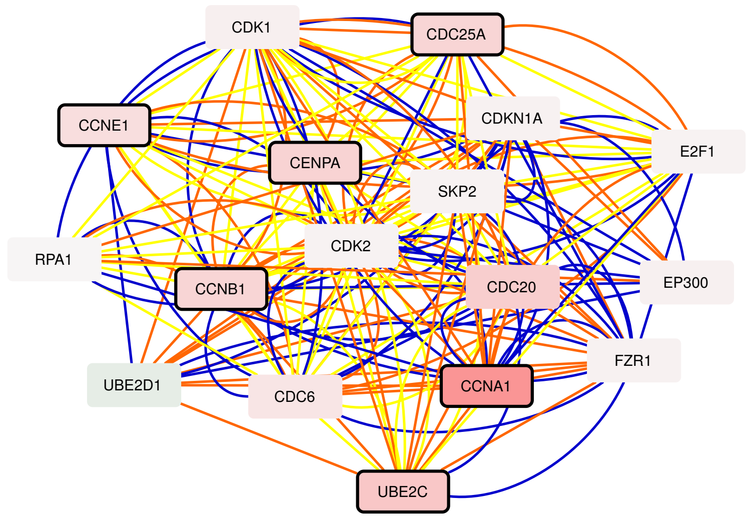


Figure S12.16. Yao's dataset, myotubes: Active module 16

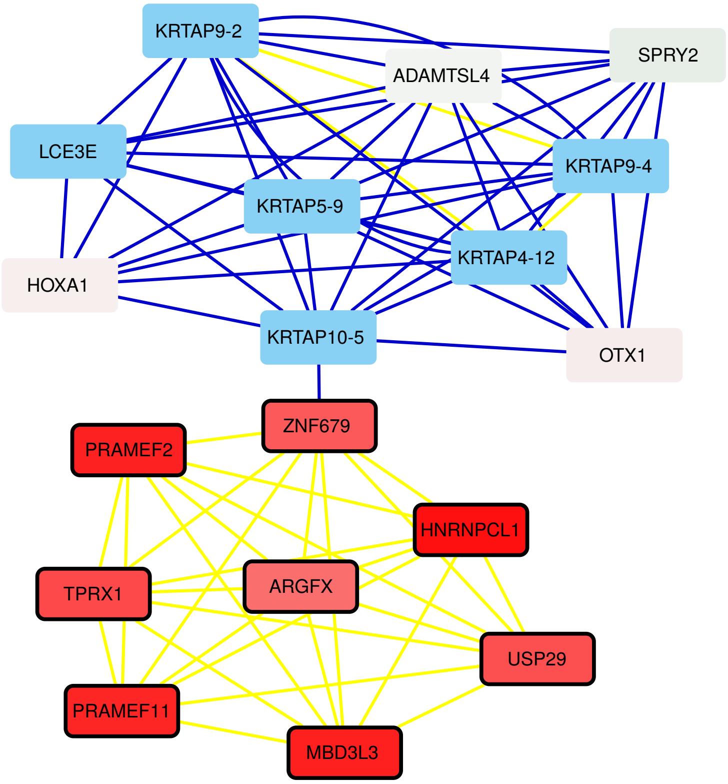


Figure S12.17. Yao's dataset, myotubes: Active module 17

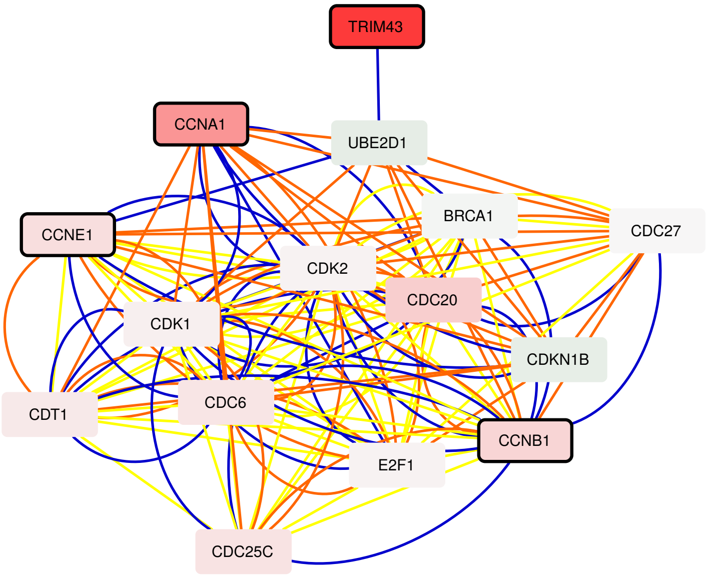


Figure S12.18. Yao's dataset, myotubes: Active module 18

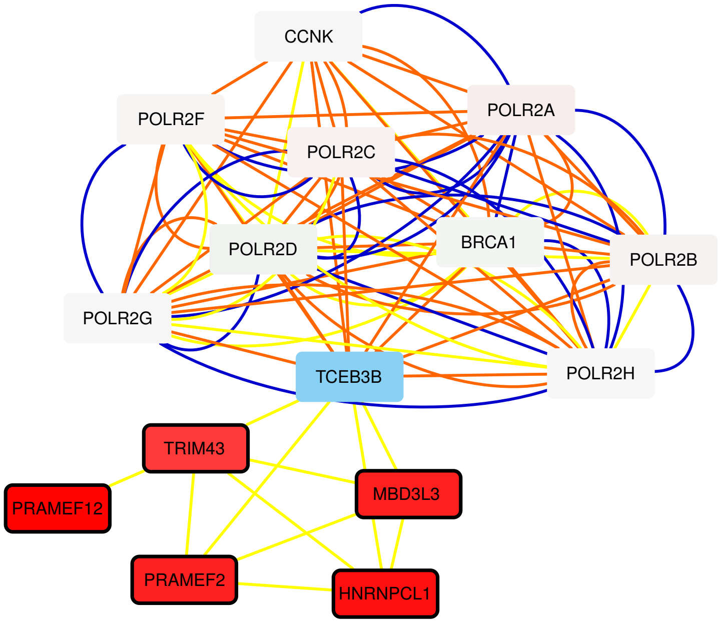


Figure S12.19. Yao's dataset, myotubes: Active module 19

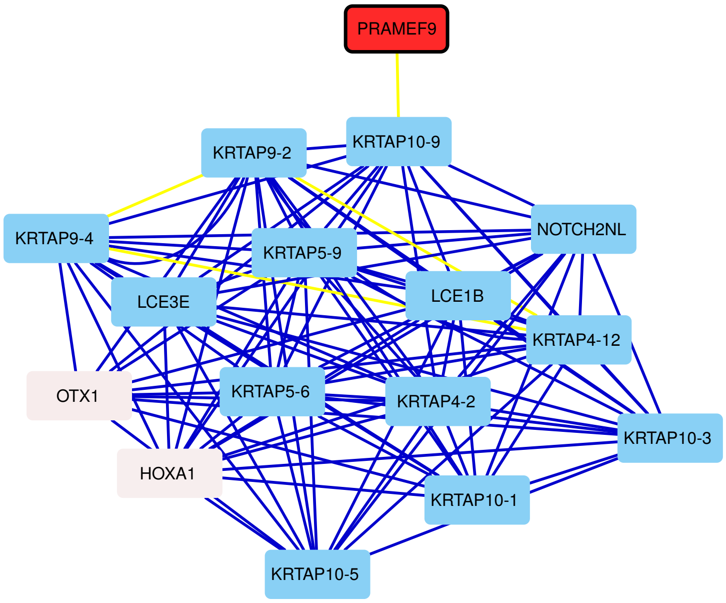


Figure S12.20. Yao's dataset, myotubes: Active module 20

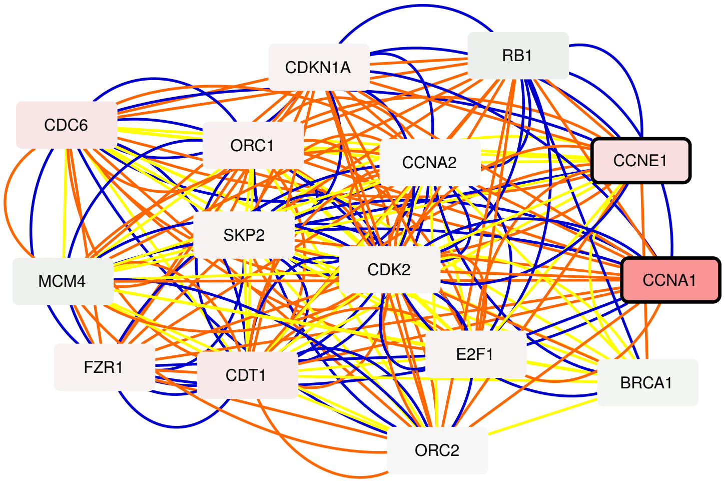


Figure S12.21. Yao's dataset, myotubes: Active module 21

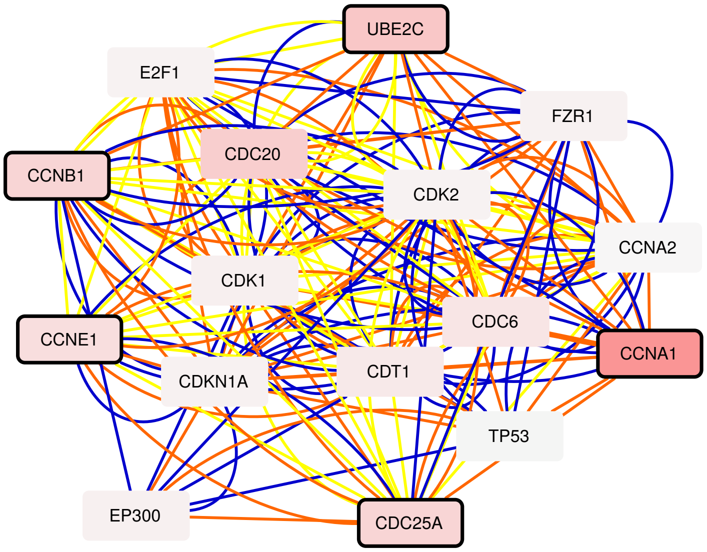


Figure S12.22. Yao's dataset, myotubes: Active module 22

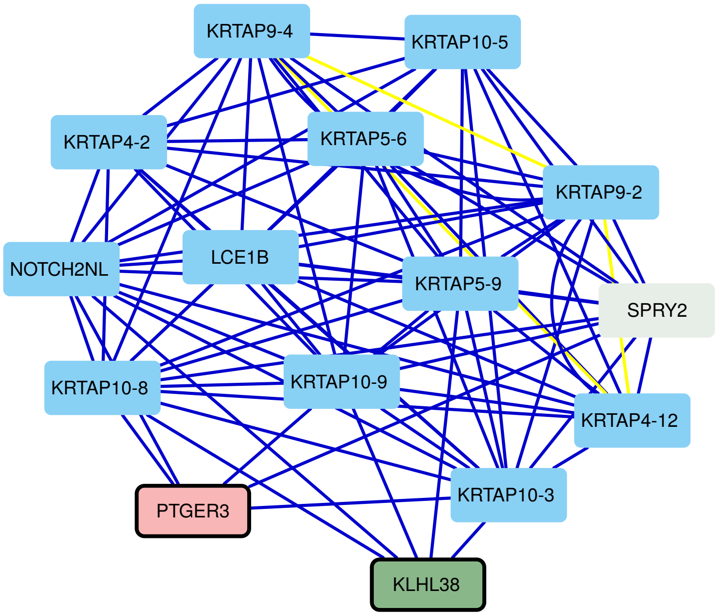


Figure S12.23. Yao's dataset, myotubes: Active module 23

#### Figure S13

Twenty three active modules obtained by applying MOGAMUN on Banerji's 2017 dataset [2] (see Table S2 for the list of samples). The color of the nodes represents the fold-change, where green and red nodes correspond to under- and over-expressed genes, respectively. Nodes with bold black border correspond to genes significantly differentially expressed ($FDR < 0.05$ and absolute $log_2$ fold-change $> 1$). Blue and white nodes correspond to genes with no associated transcriptomics data and no deregulation, respectively.

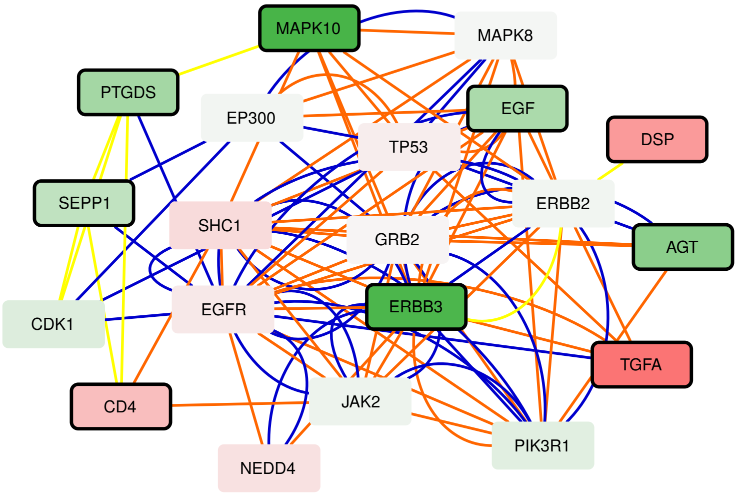


Figure S13.1. Banerji's 2017 dataset: Active module 1

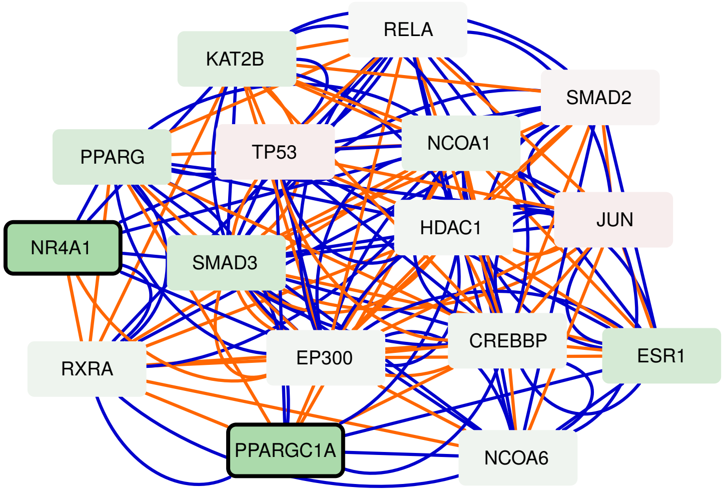


Figure S13.2. Banerji's 2017 dataset: Active module 2

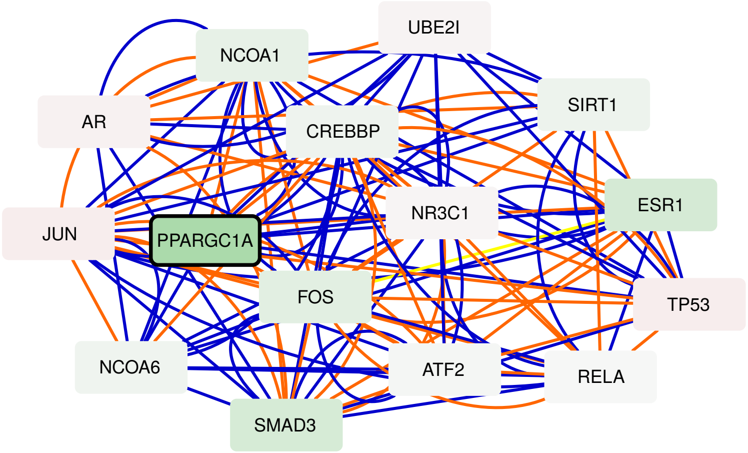


Figure S13.3. Banerji's 2017 dataset: Active module 3

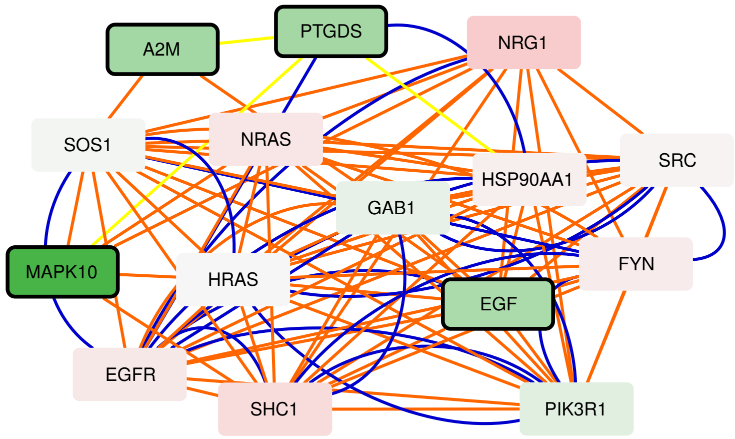


Figure S13.4. Banerji's 2017 dataset: Active module 4

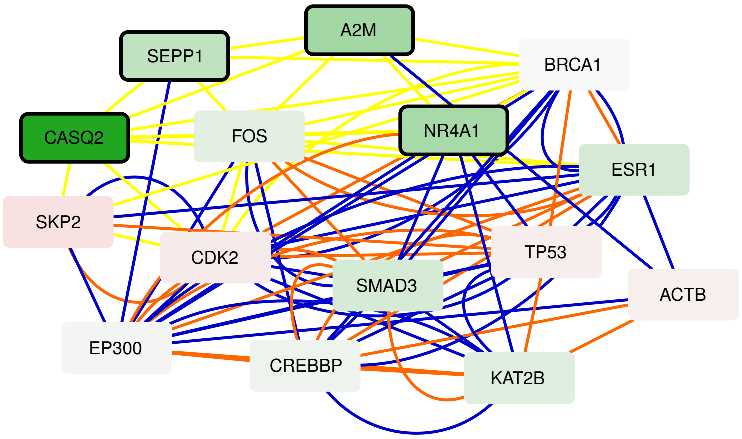


Figure S13.5. Banerji's 2017 dataset: Active module 5

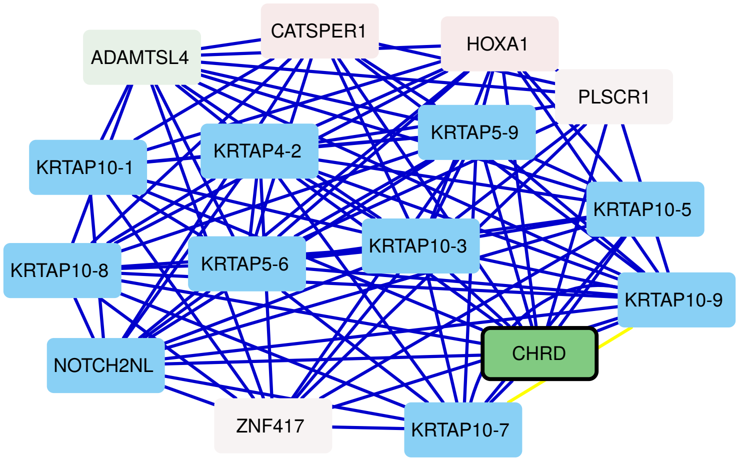


Figure S13.6. Banerji's 2017 dataset: Active module 6

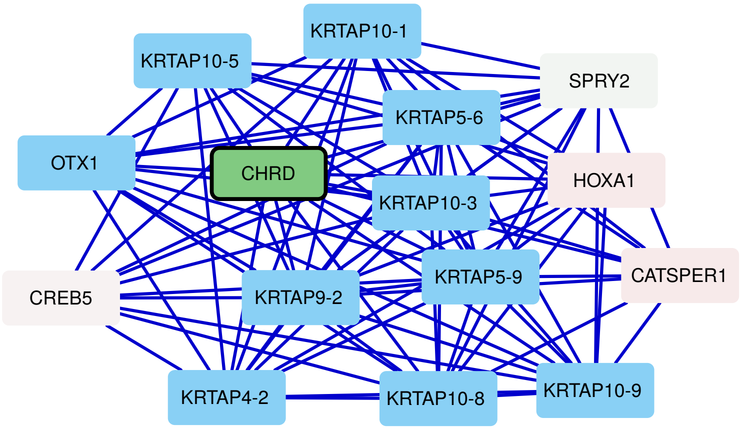


Figure S13.7. Banerji's 2017 dataset: Active module 7

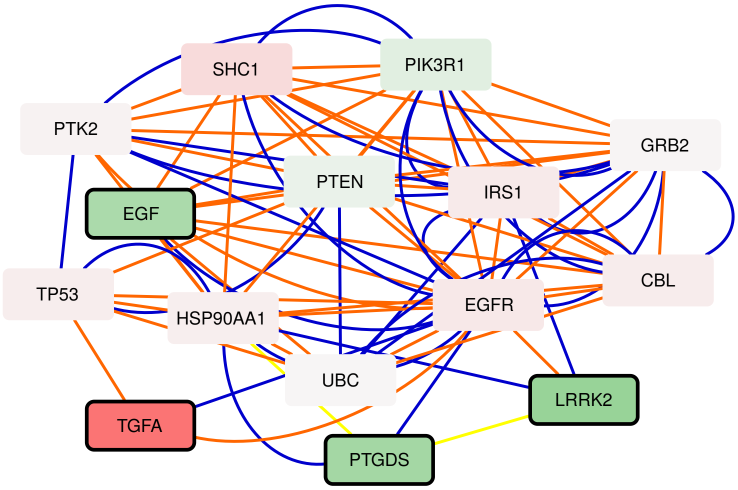


Figure S13.8. Banerji's 2017 dataset: Active module 8

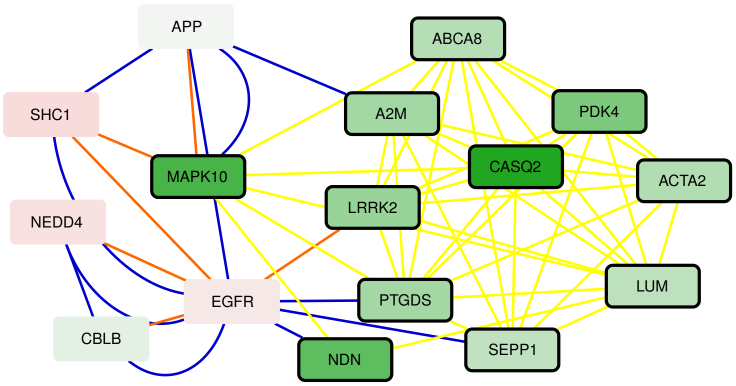


Figure S13.9. Banerji's 2017 dataset: Active module 9

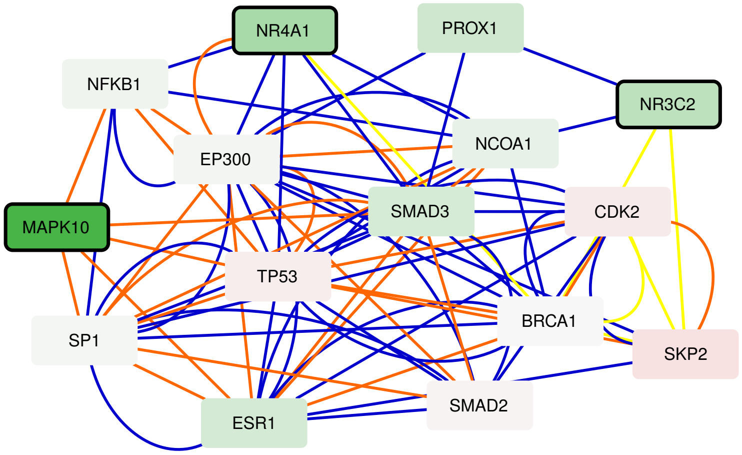


Figure S13.10. Banerji's 2017 dataset: Active module 10

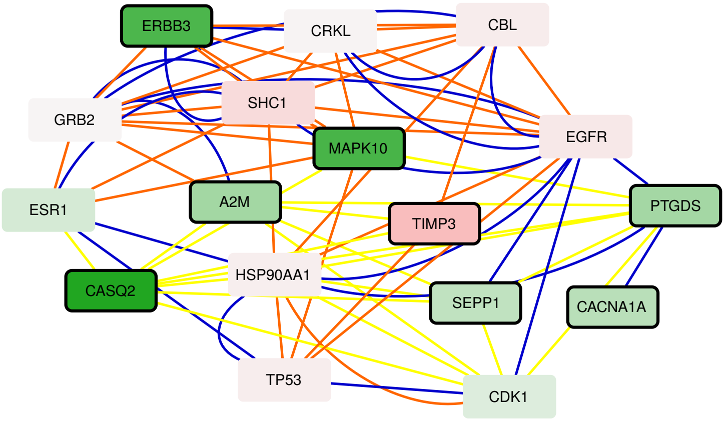


Figure S13.11. Banerji's 2017 dataset: Active module 11

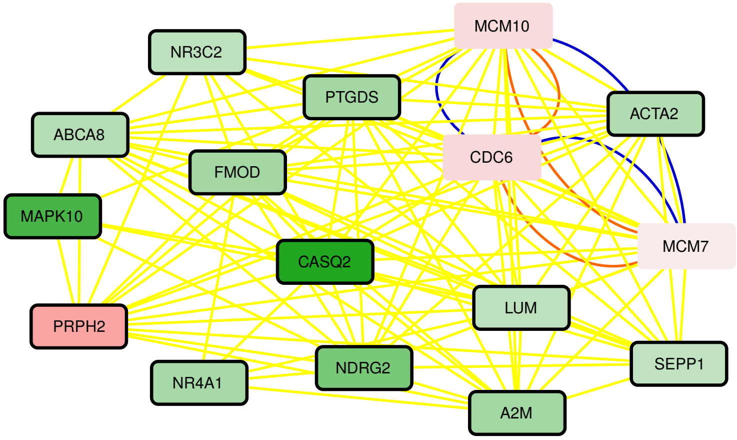


Figure S13.12. Banerji's 2017 dataset: Active module 12

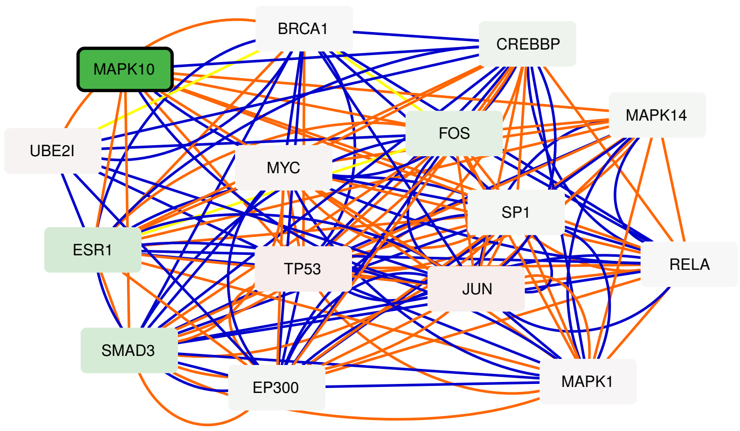


Figure S13.13. Banerji's 2017 dataset: Active module 13

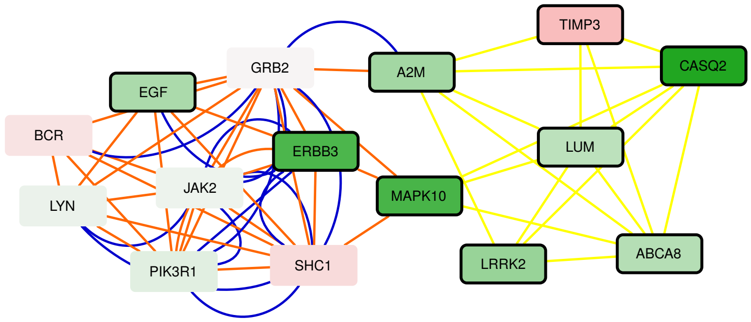


Figure S13.14. Banerji's 2017 dataset: Active module 14

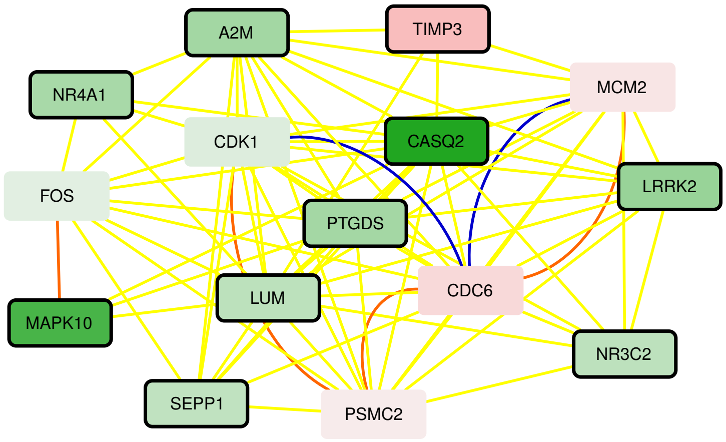


Figure S13.15. Banerji's 2017 dataset: Active module 15

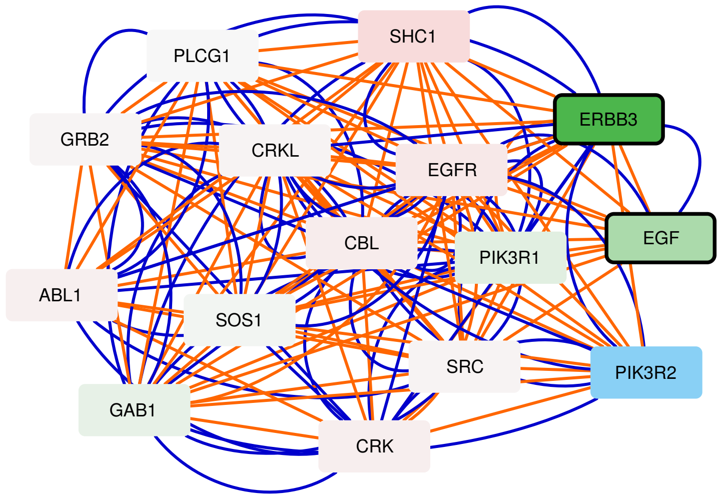


Figure S13.16. Banerji's 2017 dataset: Active module 16

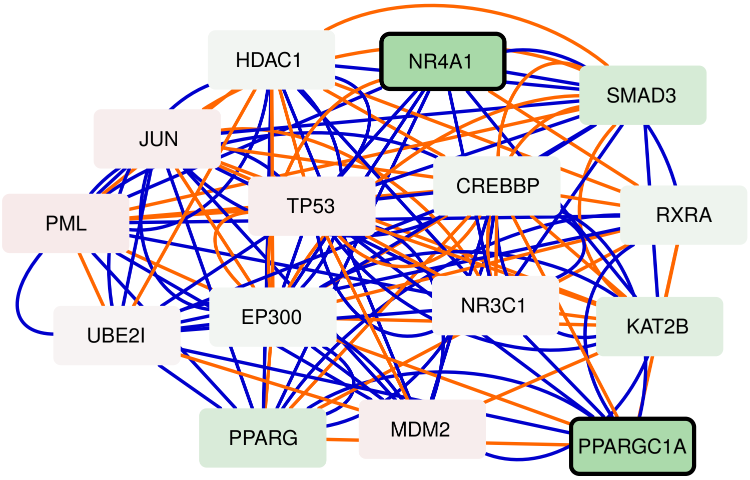


Figure S13.17. Banerji's 2017 dataset: Active module 17

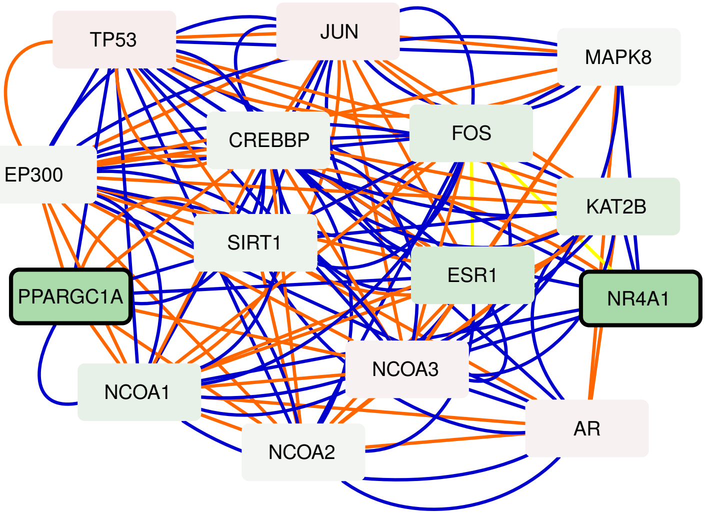


Figure S13.18. Banerji's 2017 dataset: Active module 18

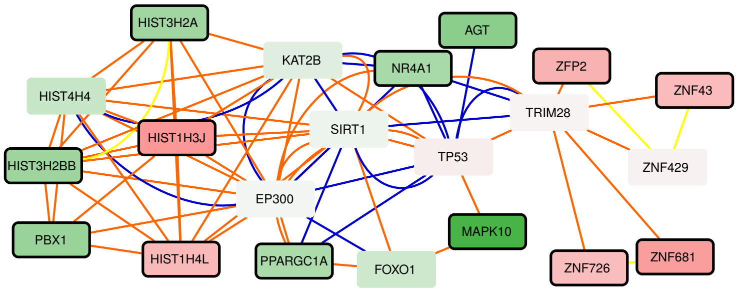


Figure S13.19. Banerji's 2017 dataset: Active module 19

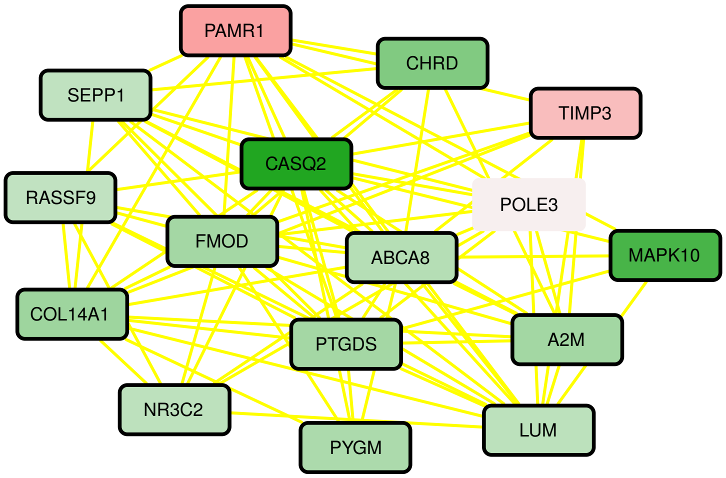


Figure S13.20. Banerji's 2017 dataset: Active module 20

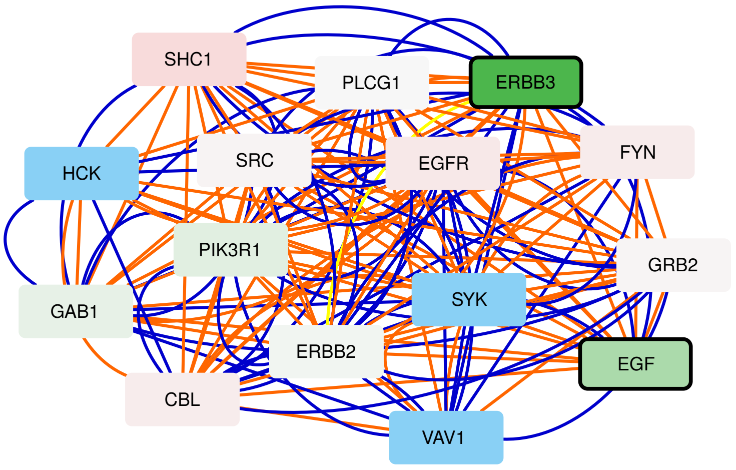


Figure S13.21. Banerji's 2017 dataset: Active module 21

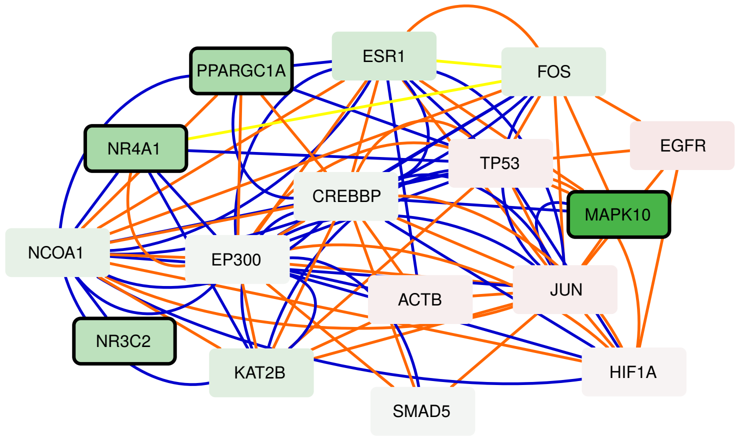


Figure S13.22. Banerji's 2017 dataset: Active module 22

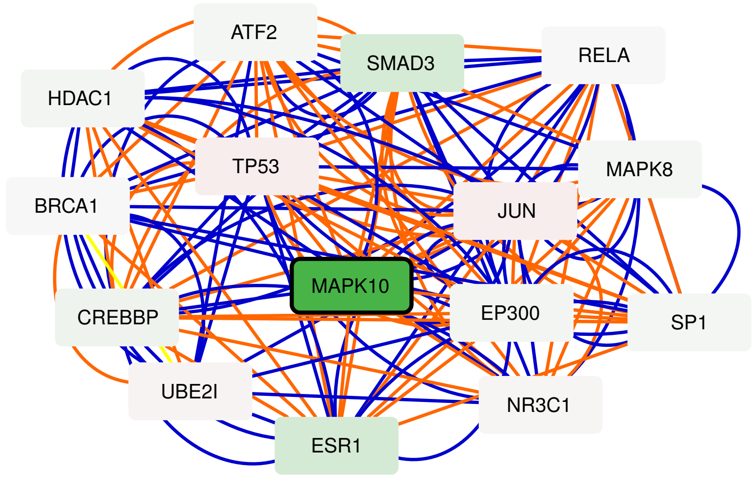


Figure S13.23. Banerji's 2017 dataset: Active module 23

### Figure S14

Seventeen active modules obtained by applying MOGAMUN on Banerji's 2019 dataset [3] (see Table S3 for the list of samples). The color of the nodes represents the fold-change, where green and red nodes correspond to under- and over-expressed genes, respectively. Nodes with bold black border correspond to genes significantly differentially expressed ($FDR < 0.05$ and absolute $log_2$ fold-change $> 1$). Blue and white nodes correspond to genes with no associated transcriptomics data and no deregulation, respectively.

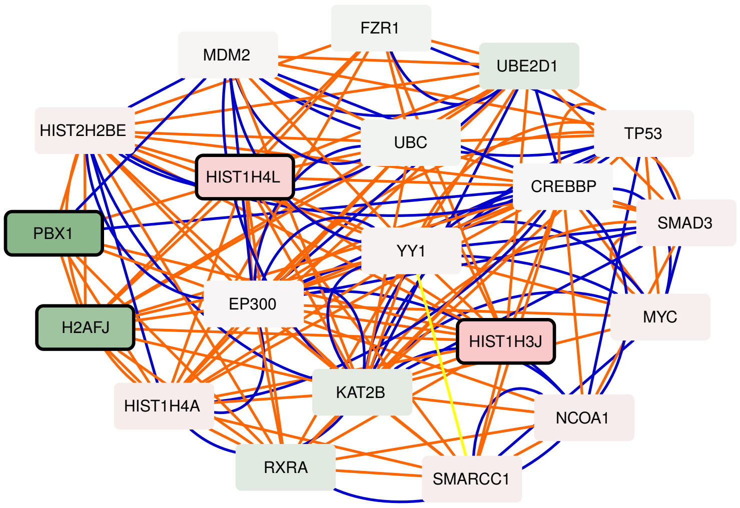


Figure S14.1. Banerji's 2019 dataset: Active module 1

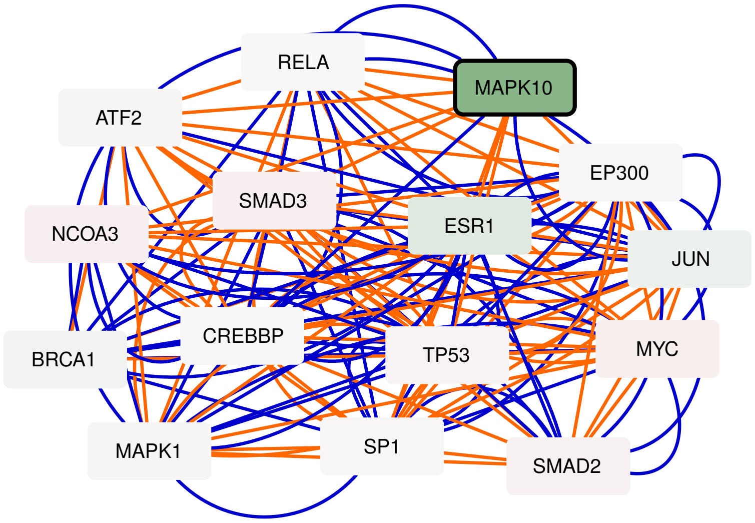


Figure S14.2. Banerji's 2019 dataset: Active module 2

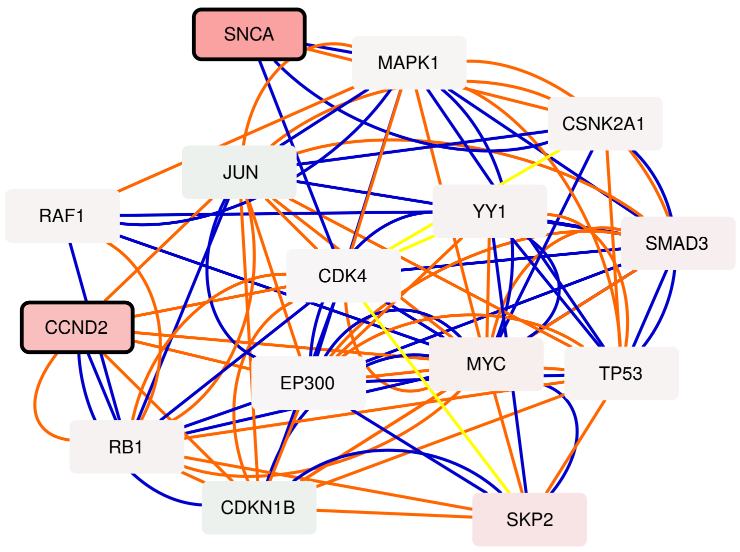


Figure S14.3. Banerji's 2019 dataset: Active module 3

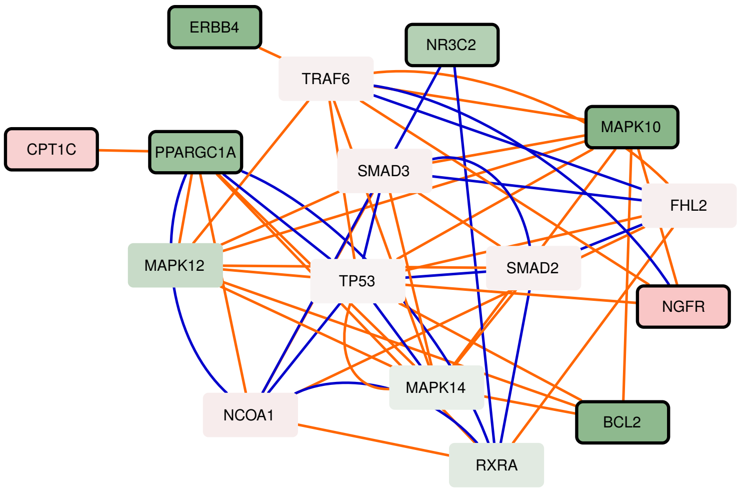


Figure S14.4. Banerji's 2019 dataset: Active module 4

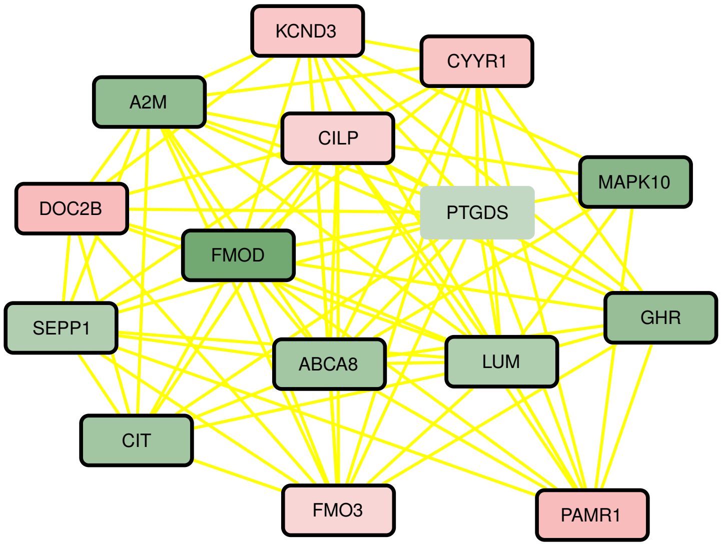


Figure S14.5. Banerji's 2019 dataset: Active module 5

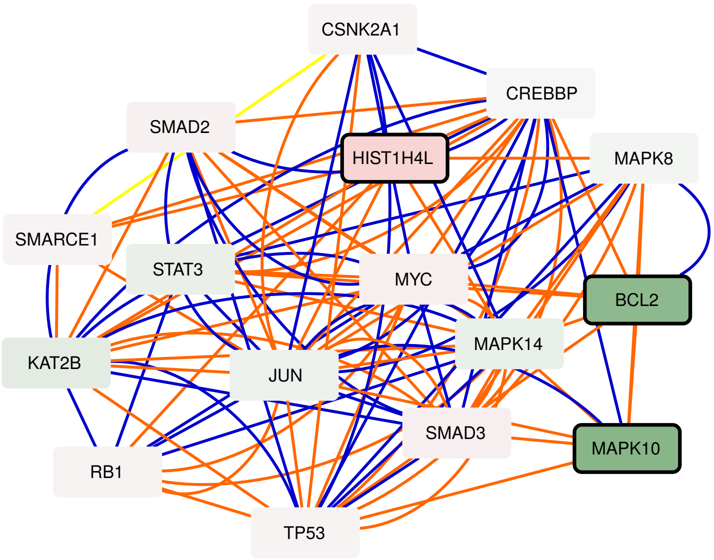


Figure S14.6. Banerji's 2019 dataset: Active module 6

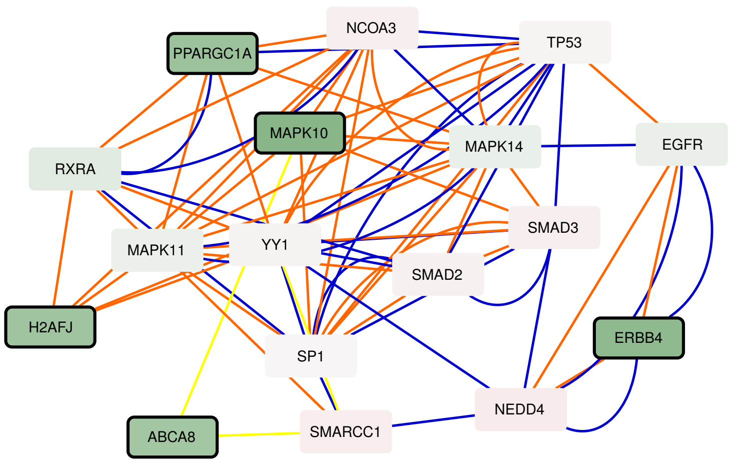


Figure S14.7. Banerji's 2019 dataset: Active module 7

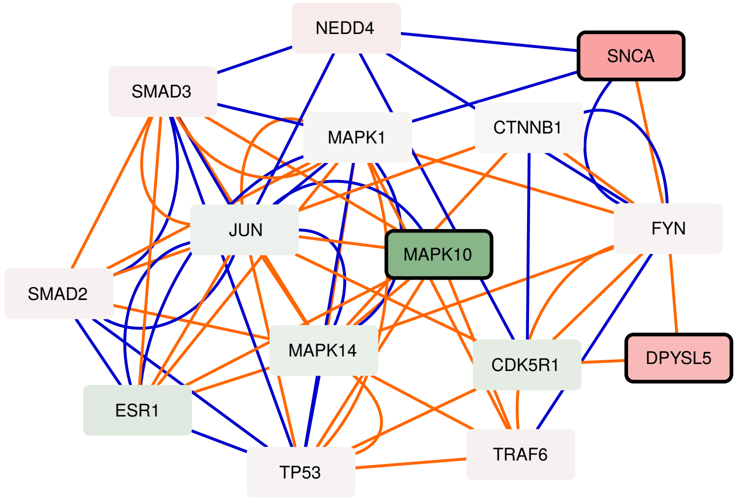


Figure S14.8. Banerji's 2019 dataset: Active module 8

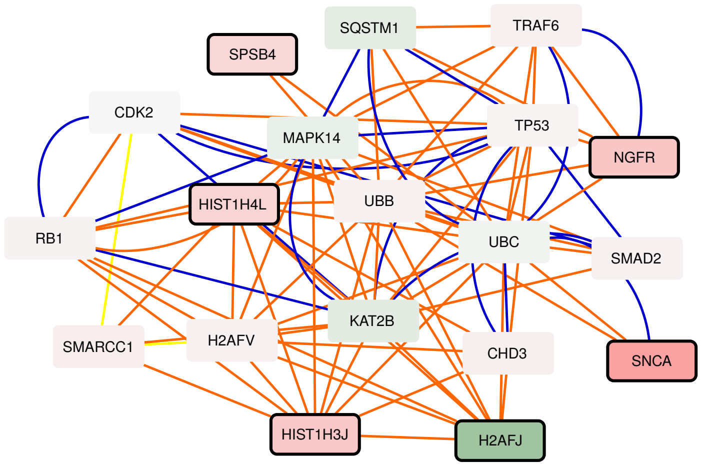


Figure S14.9. Banerji's 2019 dataset: Active module 9

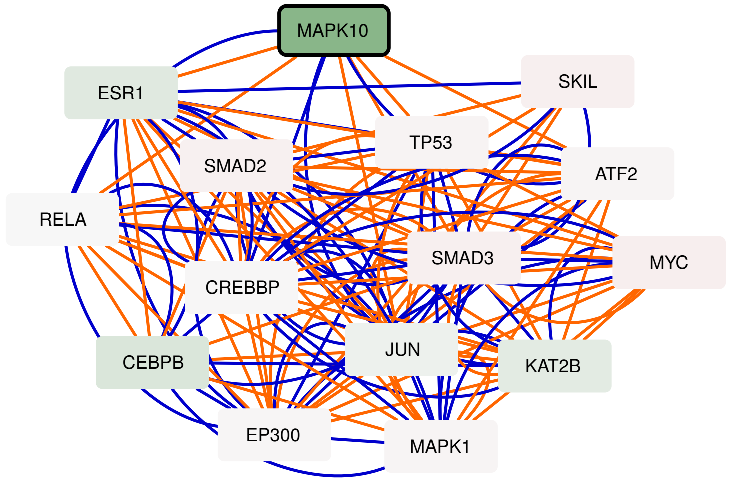


Figure S14.10. Banerji's 2019 dataset: Active module 10

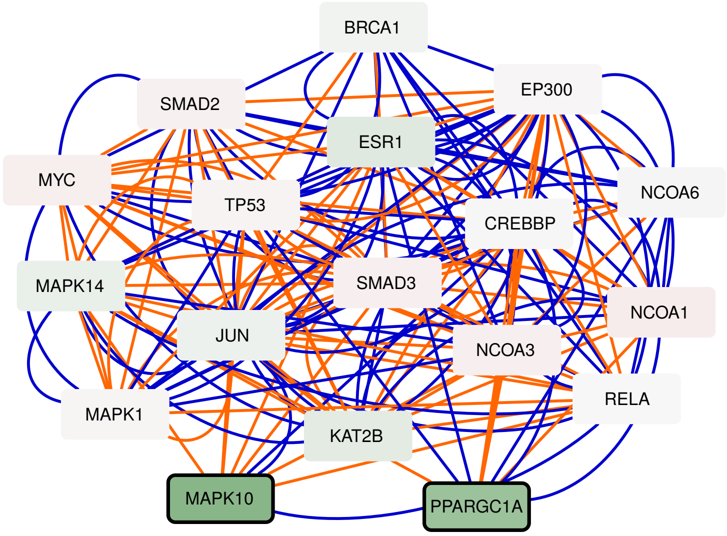


Figure S14.11. Banerji's 2019 dataset: Active module 11

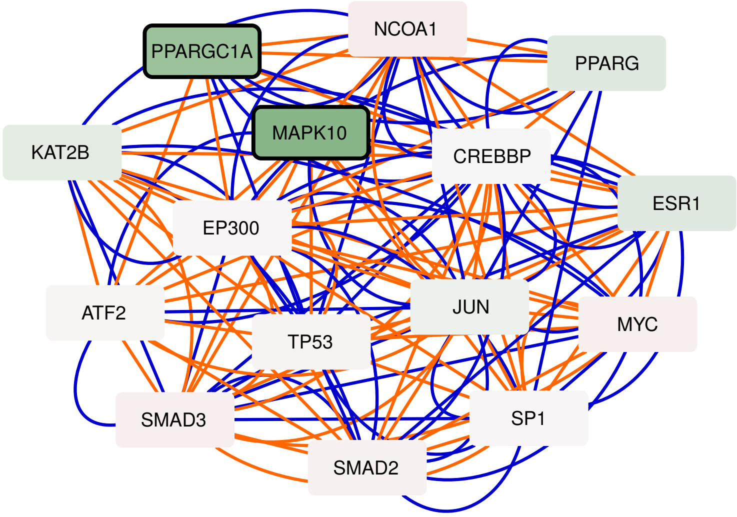


Figure S14.12. Banerji's 2019 dataset: Active module 12

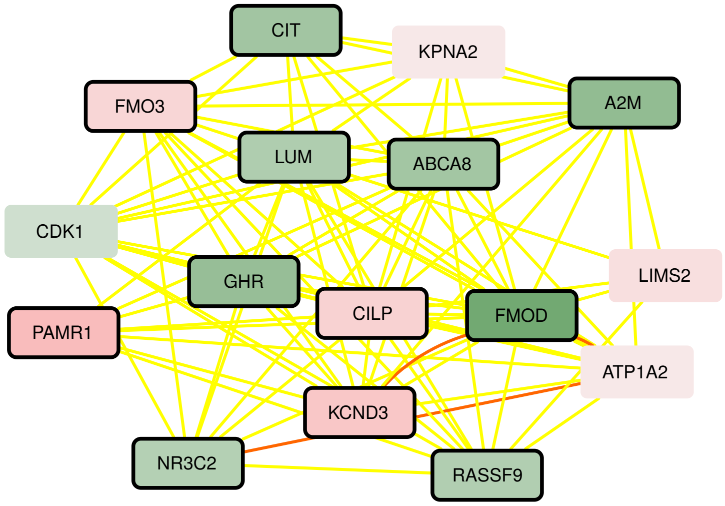


Figure S14.13. Banerji's 2019 dataset: Active module 13

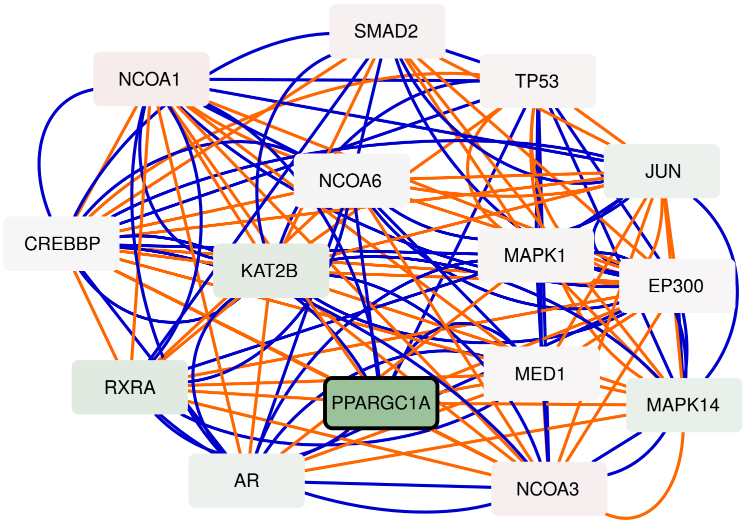


Figure S14.14. Banerji's 2019 dataset: Active module 14

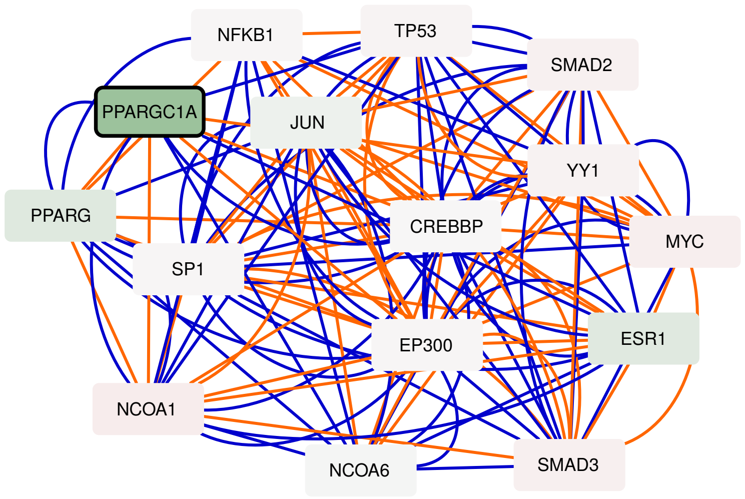


Figure S14.15. Banerji's 2019 dataset: Active module 15

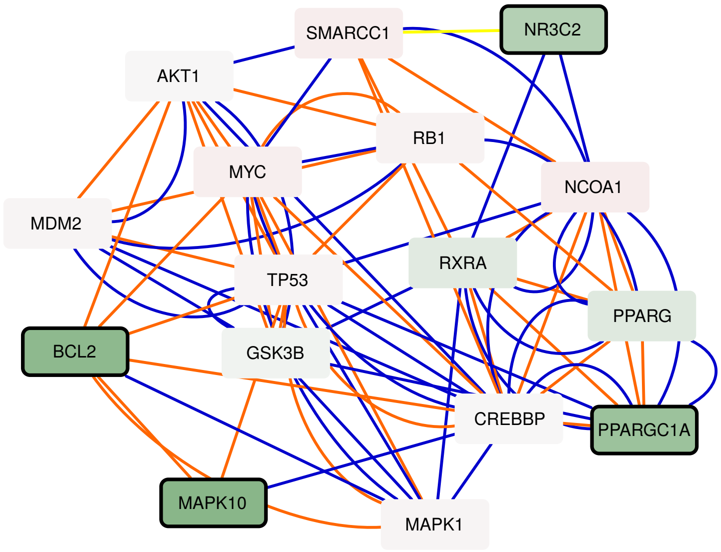


Figure S14.16. Banerji's 2019 dataset: Active module 16

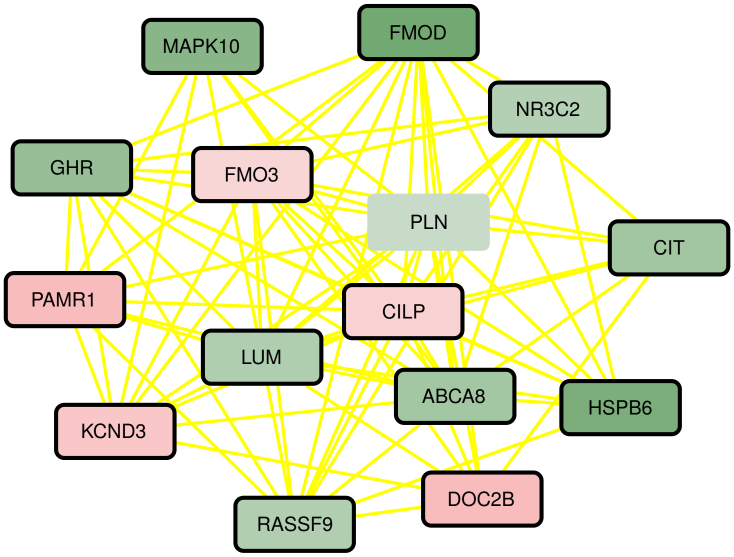


Figure S14.17. Banerji's 2019 dataset: Active module 17

| Sample_ID | Type | Origin |
|---|---|---|
| F1 | Patient | Biopsy |
| F2 | Patient | Biopsy |
| F3 | Patient | Biopsy |
| F4 | Patient | Biopsy |
| F5 | Patient | Biopsy |
| F6 | Patient | Biopsy |
| F7 | Patient | Biopsy |
| F8 | Patient | Biopsy |
| F9 | Patient | Biopsy |
| C1 | Control | Biopsy |
| C2 | Control | Biopsy |
| C3 | Control | Biopsy |
| C4 | Control | Biopsy |
| C5 | Control | Biopsy |
| C6 | Control | Biopsy |
| C7 | Control | Biopsy |
| C8 | Control | Biopsy |
| C9 | Control | Biopsy |
| F4 | Patient | Myoblast |
| F6 | Patient | Myoblast |
| C21 | Control | Myoblast |
| C22 | Control | Myoblast |
| F4 | Patient | Myotube |
| F6 | Patient | Myotube |
| C20 | Control | Myotube |
| C21 | Control | Myotube |
| C22 | Control | Myotube |

Table S1: Samples from Yao's datasets [1]. Downloaded from https://www.ncbi.nlm.nih.gov/geo/query/acc.cgi?acc=GSE56787

| ID | Type | Batch |
|---|---|---|
| 54_12_r1 | Patient | 1 |
| 54_12_r2 | Patient | 1 |
| 54_12_r3 | Patient | 1 |
| 54_6_r1 | Control | 1 |
| 54_6_r2 | Control | 1 |
| 54_6_r3 | Control | 1 |
| 54_2_r1 | Patient | 2 |
| 54_2_r2 | Patient | 2 |
| 54_2_r3 | Patient | 2 |
| 54_A10_r1 | Control | 2 |
| 54_A10_r2 | Control | 2 |
| 54_A10_r3 | Control | 2 |
| 54_A5_r1 | Patient | 2 |
| 54_A5_r2 | Patient | 2 |
| 54_A5_r3 | Patient | 2 |
| 12ABic_r3 | Patient | 3 |
| 12ABic_r1 | Patient | 3 |
| 12ABic_r2 | Patient | 3 |
| 12UBic_r3 | Control | 3 |
| 12UBic_r2 | Control | 3 |
| 12UBic_r1 | Control | 3 |
| 16ABic_r1 | Patient | 3 |
| 16ABic_r2 | Patient | 3 |
| 16ABic_r3 | Patient | 3 |
| 16UBic_r3 | Control | 3 |
| 16UBic_r2 | Control | 3 |
| 16UBic_r1 | Control | 3 |

Table S2: Samples from Banerji's 2017 dataset [2]. Downloaded from https://www.ncbi.nlm.nih.gov/geo/query/acc.cgi?acc=GSE102812

| ID | Type | Batch |
|---|---|---|
| 54_12_T8_r1 | Patient | 1 |
| 54_12_T8_r2 | Patient | 1 |
| 54_12_T8_r3 | Patient | 1 |
| 54_6_T8_r1 | Control | 1 |
| 54_6_T8_r2 | Control | 1 |
| 54_6_T8_r3 | Control | 1 |
| 54_2_T8_r1 | Patient | 2 |
| 54_2_T8_r2 | Patient | 2 |
| 54_2_T8_r3 | Patient | 2 |
| 54_A10_T8_r1 | Control | 2 |
| 54_A10_T8_r2 | Control | 2 |
| 54_A10_T8_r3 | Control | 2 |
| 54_A5_T8_r1 | Patient | 2 |
| 54_A5_T8_r2 | Patient | 2 |
| 54_A5_T8_r3 | Patient | 2 |
| 12A_T8_r1 | Patient | 3 |
| 12A_T8_r2 | Patient | 3 |
| 12A_T8_r3 | Patient | 3 |
| 12U_T8_r1 | Control | 3 |
| 12U_T8_r2 | Control | 3 |
| 12U_T8_r3 | Control | 3 |
| 16A_T8_r1 | Patient | 3 |
| 16A_T8_r2 | Patient | 3 |
| 16A_T8_r3 | Patient | 3 |
| 16U_T8_r1 | Control | 3 |
| 16U_T8_r2 | Control | 3 |
| 16U_T8_r3 | Control | 3 |

Table S3: Samples from Banerji's 2019 dataset [3]. Downloaded from https://www.ncbi.nlm.nih.gov/geo/query/acc.cgi?acc=GSE123468

### References


[1] Z. Yao, L. Snider, J. Balog, R. J. Lemmers, S. M. Van Der Maarel, R. Tawil, and S. J. Tapscott. Dux4-induced gene expression is the major molecular signature in fshd skeletal muscle. *Human molecular genetics 23*, 2014.

[2] C. R. Banerji, M. Panamarova, H. Hebaishi, R. B. White, F. Relaix, S. Severini, and P. S. Zammit. Pax7 target genes are globally repressed in facioscapulohumeral muscular dystrophy skeletal muscle. *Nature communications 8*, 2017.

[3] C. R. Banerji, M. Panamarova, J. Pruller, N. Figeac, H. Hebaishi, E. Fidanis, ..., and P. S. Zammit. Dynamic transcriptomic analysis reveals suppression of pgc1 $\alpha$/err $\alpha$ drives perturbed myogenesis in facioscapulohumeral muscular dystrophy. *Human molecular genetics 28*, 2019.
